# Ubiquitination drives COPI priming and Golgi SNARE localization

**DOI:** 10.1101/2021.11.29.470439

**Authors:** Swapneeta S. Date, Peng Xu, Nathaniel L. Hepowit, Nicholas S. Diab, Jordan Best, Boyang Xie, Jiale Du, Eric R. Strieter, Lauren Parker Jackson, Jason A. MacGurn, Todd R. Graham

**Affiliations:** Biological Sciences, Vanderbilt University, Nashville, TN 37212; Cell and Developmental Biology, Vanderbilt University, Nashville, TN 37212; Department of Chemistry, University of Massachusetts Amherst, Amherst, MA 01003, USA

## Abstract

Deciphering mechanisms controlling SNARE localization within the Golgi complex is crucial to understanding protein trafficking patterns within the secretory pathway. SNAREs are also thought to prime COPI assembly to ensure incorporation of these essential cargoes into vesicles but the regulation of these events is poorly understood. Here we report a roles for ubiquitin recognition by COPI in SNARE trafficking and in stabilizing interactions between Arf, COPI, and Golgi SNAREs. The ability of COPI to bind ubiquitin through its N-terminal WD repeat domain of β’COP or through an unrelated ubiquitin-binding domain (UBD) is essential for the proper localization of Golgi SNAREs Bet1 and Gos1. We find that COPI, the ArfGAP Glo3 and multiple Golgi SNAREs are ubiquitinated. Notably, the binding of Arf and COPI to Gos1 is markedly enhanced by ubiquitination of these components. Glo3 is thought to prime COPI-SNARE interactions; however, Glo3 is not enriched in the ubiquitin-stabilized SNARE-Arf-COPI complex but is instead enriched with COPI complexes that lack SNAREs. These results support a new model for how posttranslational modifications drive COPI priming events crucial for Golgi SNARE localization.

## Introduction

The sorting of proteins in the endomembrane system is a highly regulated, vesicle-mediated process important for many physiological events. Coat proteins drive the formation of vesicles by assembling onto the cytosolic surface of cellular membranes, where they select cargo proteins^1–4^. COPI-coated vesicles originate at the Golgi, and mediate retrograde transport between Golgi cisternae or back to the ER ^1, 5, 6^ COPI is a highly conserved heptameric protein complex (α, β, β’, γ, δ, ε and ζ subunits) that is recruited to Golgi membranes by the small GTP-binding protein Arf (Arf1 and Arf2 in budding yeast)^7–10^. The N-terminal WD repeat (WDR) domains of α- and β’-COP recognize sorting signals on cargoes, such as dilysine motifs commonly found on ER-resident membrane proteins^11–13^. As Golgi cisternae mature from *cis* to *trans* in budding yeast, the retrograde movement of resident proteins becomes critical in order to maintain a functional Golgi. Resident Golgi proteins, thus, are also important COPI cargo^14–17^.

SNAREs are another critical cargo of COPI vesicles because they are essential for vesicle fusion with the target membrane and are proposed to prime, or nucleate, coat formation^18–21^. In addition to incorporating v-SNAREs into vesicle membranes, COPI must also mediate retrograde transport of early Golgi t-SNAREs moving through the Golgi by cisternal maturation to maintain Golgi organization, but how COPI mediates sorting of SNAREs is poorly understood. Because of the tail-anchored topology of SNAREs, none of these proteins contain a C-terminal dilysine motif on the cytosolic side of the membrane where it is accessible to COPI. Few sorting signals have been identified in SNARE proteins and how they are incorporated into COPI vesicles is incompletely understood^22–29^. The ArfGAP protein Glo3 may contribute because it is known to interact with COPI, Arf-GTP and SNAREs and is proposed to be part of the priming complex^19^. However, Glo3 stimulates GTP hydrolysis by Arf to form Arf-GDP, which destabilizes the COPI coat and is crucial for vesicle uncoating^30^. How these Glo3 interactions are regulated to allow Arf-GTP mediated COPI assembly during formation and their influence on SNARE localization is unclear.

We recently found that COPI plays a role in the recycling of a budding yeast v-SNARE, Snc1, from the endocytic pathway to the TGN through recognition of a polyubiquitin (polyUb) signal^31^. The cargo-binding WDR domains of α-COP and β’-COP bind specifically to polyUb^31^. Deletion of the β’-COP N-terminal WDR domain (β’-COP Δ2-304) disrupts Snc1 recycling while replacement of this domain with unrelated ubiquitin-binding domains restores Snc1 recycling. Thus, β’-COP plays a critical role in this ubiquitin-dependent trafficking route, but it was unclear if the COPI-ubiquitin interaction is important for trafficking of any other cargoes. In the current study, we seek to determine if the COPI-ubiquitin interaction is a general principle of SNARE trafficking. We show that the normal localization of several SNAREs functioning at the ER-Golgi interface or within the Golgi, including Bet1, Gos1, Snc2, Bos1 and Sec22, requires COPI-ubiquitin interactions. In addition, several Golgi SNAREs, COPI subunits, and the ArfGAP Glo3 are ubiquitinated with non-degradative ubiquitin linkages under physiological conditions in *Saccharomyces cerevisiae*. Importantly, we show that ubiquitination of these components strengthens the interaction between Golgi SNAREs and COPI while apparently excluding Glo3, providing critical new mechanistic insight into potential priming mechanisms for COPI vesicle formation. These studies highlight the finely orchestrated role of posttranslational modification in driving COPI priming and sorting of a specific set of Golgi SNAREs crucial to the functional organization of Golgi.

## Results

### SNAREs mislocalize to morphologically aberrant compartments in the β’-COP Δ2-304 mutant

To determine the dependence of SNARE localization on COPI-ubiquitin interactions, we individually tagged 16 yeast SNAREs with mNeonGreen (mNG) and expressed them in *S. cerevisiae* wild-type (WT) cells or in a strain where the ubiquitin-binding N-terminal WDR of β’-COP had been deleted (β’-COP Δ2-304) (**Fig. 1a and Extended Data Table 1**). This β’-COP mutation does not completely eliminate COPI polyUb binding because α-COP can also bind polyUb; therefore, the SNAREs were overexpressed from a strong ADH promoter so the screen would be more sensitive for detecting changes in localization^31^. Many of the mNG-SNARE fluorescent patterns were indistinguishable between WT and β’-COP Δ2-304 cells (**Extended Data Fig. 1a**). However, for Bet1, Gos1, Snc1, and Snc2, a significant accumulation of individual SNAREs to elongated tube-like structures and ring-like structures was observed in β’-COP Δ2-304 **(Fig. 1b-c)**. As previously shown for Snc1^31^, Snc2 plasma membrane localization was also reduced in the COPI mutant. In addition, Sec22 and Bos1 were partially mislocalized to vacuolar structures in β’-COP Δ2-304 cells (**Extended Data Fig. 1b)**. GFP is typically cleaved from protein chimeras upon arrival in the vacuole. Consistently, immunoblotting of cell lysates with anti-GFP indicated that 40% of the GFP-Sec22 chimera was cleaved in β’-COP Δ2-304 cells to release free GFP, in contrast to WT cells where less than 5% was cleaved **(Extended Data Fig. 1c-d)**.

**Figure 1.**
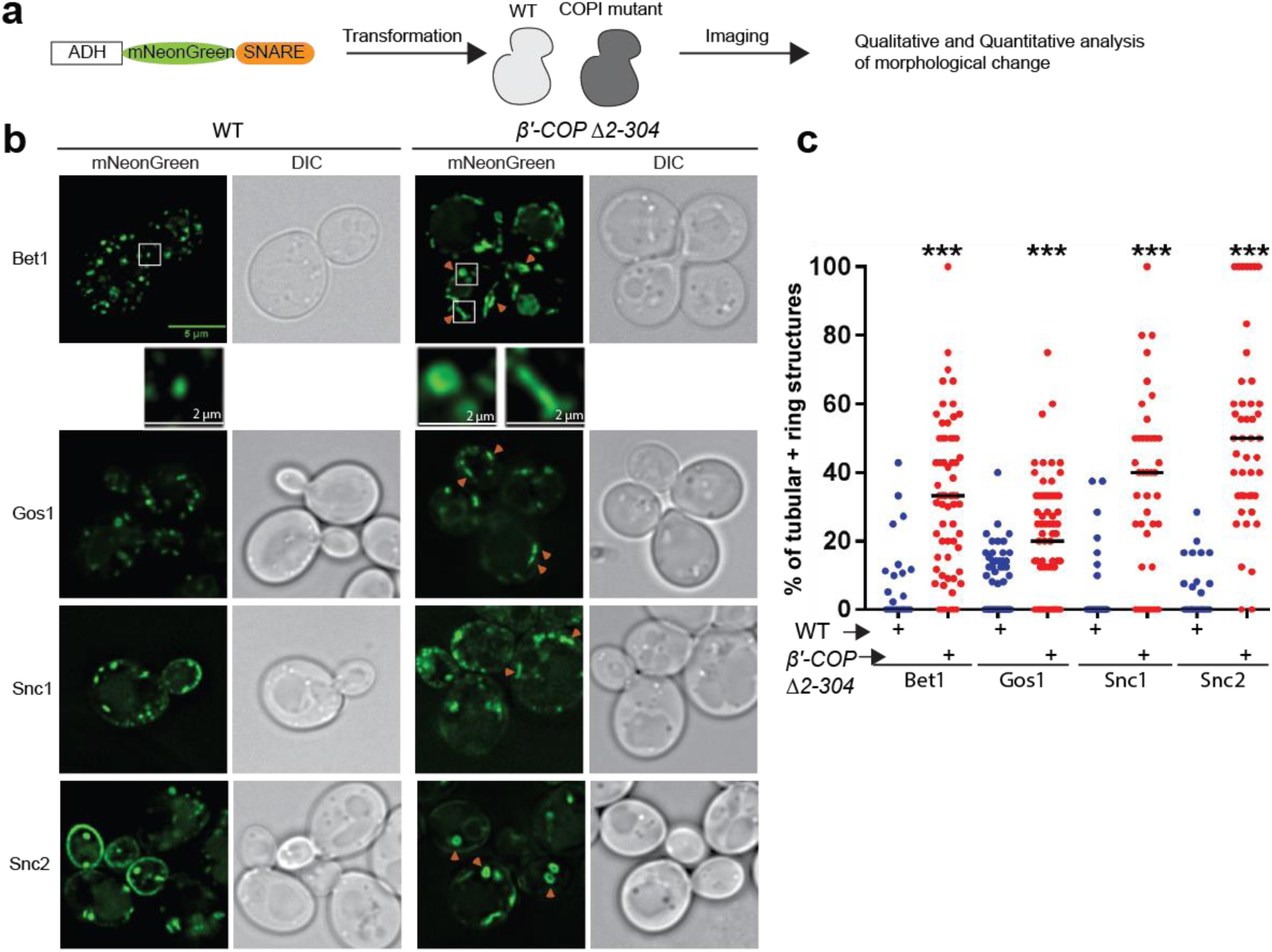
SNAREs localize to morphologically aberrant structures in COPI mutants: **(a)** Schematics of the experimental setup wherein SNAREs are individually tagged with mNeonGreen and expressed under constitutive ADH promoter in *Saccharomyces cerevisiae* wild-type (WT) cells or in cells with deleted N-terminal WDR of β’-COP (Δ2-304). **(b-c)** Panels show live-cell imaging data in *Saccharomyces cerevisiae*. Significant morphological changes are observed for SNAREs Bet1, Gos1, Snc1 and Snc2 wherein the elongated tube-like structures and ring structures (orange arrowheads) are seen in β’COP Δ2-304 cells compared to control cells with full-length β’COP (WT). Statistical differences were determined using a one-way ANOVA on the means of the three biological replicates (***p<0.001). Scale bar in represents 5 µm.

To test whether the observed morphological changes were caused by SNARE overexpression or loss of the β’-COP WDR domain, we expressed the Bet1, Snc1, and Snc2 mNG constructs using the weaker, inducible *CUP1* promoter with a short (1 hr) induction time to approximate physiological protein abundance^32^. Comparable morphological changes were observed with these SNAREs localizing to elongated tubular and ring-like structures in β’-COP Δ2-304 relative to WT cells **(Extended Data Fig. 2a-b)**. Thus, Bet1, Snc1, and Snc2 were localized to aberrant structures whether they were expressed using a strong, constitutive ADH promoter or the weaker, inducible *CUP1* promoter. All subsequent imaging studies used the *CUP1* promoter to drive SNARE expression. Together, these data suggest a dependence of a subset of SNAREs on COPI-ubiquitin interactions for their proper localization.

To further characterize the morphological changes observed for SNAREs, we performed colocalization analysis of Bet1 with Golgi markers. Bet1 significantly colocalized with medial-to late-Golgi marker Aur1 and did not significantly colocalize with the cis-Golgi marker Sed5 or TGN marker Sec7 in WT cells **(Extended Data Fig. 3a-b)**. In β’-COP Δ2-304, Bet1 is observed in elongated tubular and ring-like structures **(Fig. 1b, 2c)**, but no apparent changes were observed in the morphology of Golgi membranes containing Sed5, Aur1, and Sec7 in β’-COP Δ2-304 relative to WT cells (**Extended data Fig. 4).**

**Figure 2.**
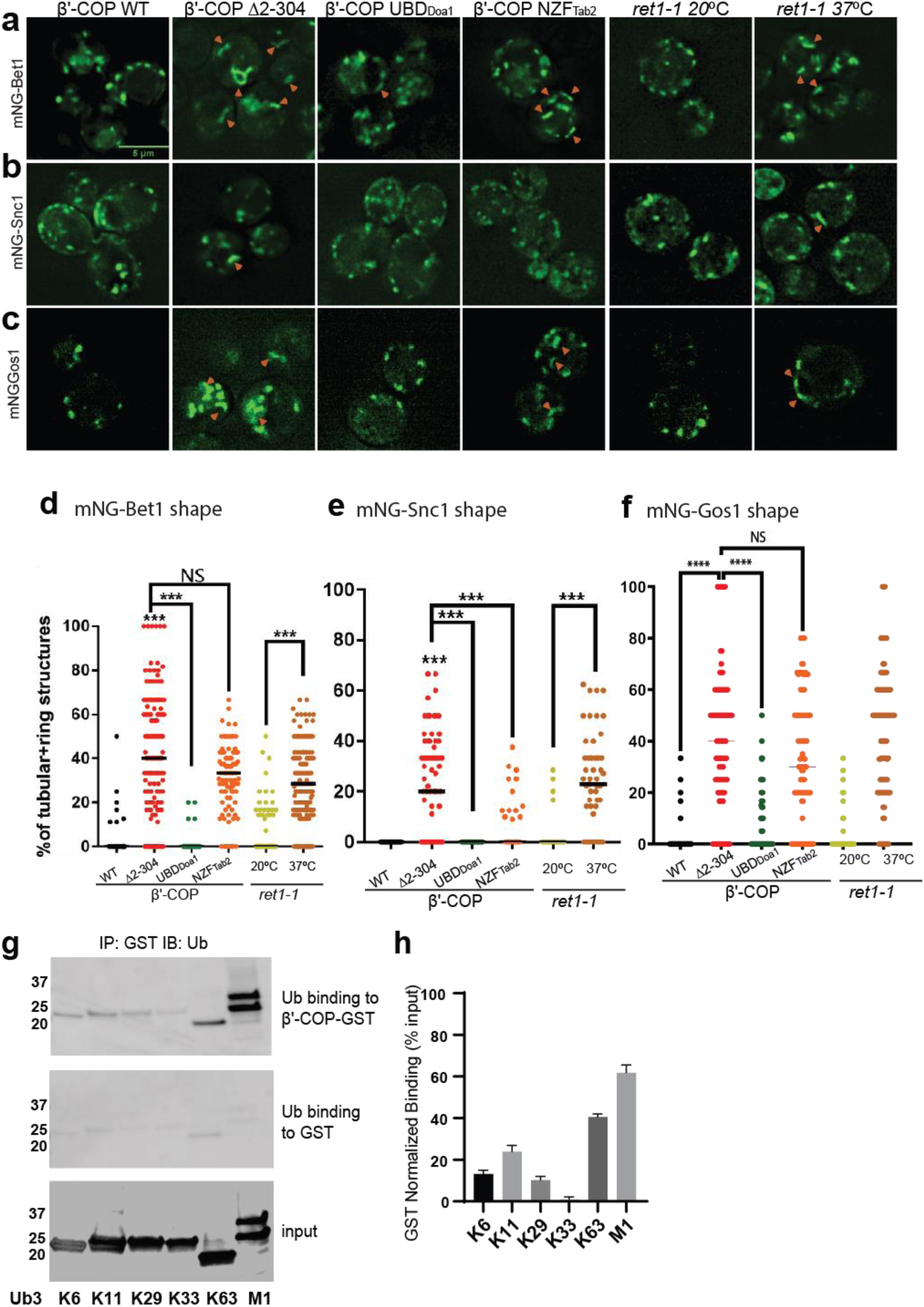
β’COP binding to ubiquitin is critical for proper SNARE localization. **(a-c)** Deletion of the N-terminal ubiquitin-binding WDR of β’COP (*Δ*2-304) leads to mislocalization of **(a)** mNG-Bet1, **(b)** mNG-Snc1 and **(c)** mNG-Gos1 into elongated tubular and ring-like structures (orange arrowheads). This phenotype is rescued by the replacement of the N-terminal ubiquitin-binding WDR of β’COP by the general ubiquitin bunding domain Doa1 (β’-COP UBD_Doa1_). The replacement of N-terminal UBD of β’COP with K63-specific UBD, NZF_Tab2_ (β’COP NZF_Tab2_) rescues the mislocalization phenotype for Snc1 but not for Bet1. The mislocalization of Bet1 and Snc1 observed in β’COP Δ2-304 cells comparable to COPI inactivation phenotype observed for *ret1-1* at nonpermissive temperatures. **(d-f)** Statistical differences were determined using a one-way ANOVA on the means of the three biological replicates (***p<0.001). **(g)** GST-β’COP (1-604) binds linear and K63-linked triUb and to some extent to K6, K11 and K29 triUb relative to the GST-only control. 0.5 mM of GST and GST tagged WDR proteins immobilized glutathione beads were incubated 250 nM Ub3 for corresponding linkages. **(h)** Quantitation of Ub3 polymers binding (GST-only background subtracted) relative to input. The values represent mean ± SEM from three independent binding experiments. Scale bar in represents 5 µm.

Whereas in WT cells Bet1 and Aur1 colocalized, Bet1 was largely mislocalized to structures in the COPI mutant that were deficient in Aur1 (**Extended Data Fig. 3c-d**). In addition, Bet1 does not traverse the plasma membrane in WT or β’-COP Δ2-304 cells and does not recycle back through the ER, consistent with a prior report^33^ **(Extended Data Fig. 3e-f)**. Thus, the COPI mutation used here does not cause whole organelle-level changes in Golgi morphology, and it appears that Bet1 is mislocalizing to a downstream (*trans-*Golgi) compartment in the COPI mutants tested **(Fig. 1b, Extended Data Fig. 3c,d)**.

### β’COP binding to ubiquitin is essential for proper SNARE localization

The N-terminal WDR domain of β’COP binds ubiquitin, and the COPI-ubiquitin interaction is critical for Snc1 retrieval^31^. To determine whether mislocalization of other SNAREs in β’-COP *Δ*2-304 is due to the inability of β’COP to bind ubiquitin, we used a set of COPI constructs **(Extended Data Fig. 5b-e)** where the N-terminal WDR domain of β’COP was replaced with (A) a general ubiquitin-binding domain of Doa1 (UBD_Doa1_), which is known to bind ubiquitin irrespective of the ubiquitin linkage type^34^ and (B) a ubiquitin-binding domain from Tab2 (NZF_tab2_) which specifically binds K63-ubiquitin linkages^35^. Compared to WT cells, Bet1, Gos1, and Snc1 were mislocalized to elongated tube structures in β’-COP Δ2-304 cells as seen previously **(Fig. 1b, Fig. 2a-f)**. Replacement of the β’-2-304 domain with the general ubiquitin-binding domain UBD_Doa1_ restored SNARE localization to punctate structures comparable to WT cells **(Fig. 2a-f)**. Surprisingly, however, the K63-linkage restricted β’COP-NZF_Tab2_ construct did not significantly correct the Bet1 or Gos1 localization pattern. We previously found that the β’COP Δ2-304 Snc1 recycling defect was fully corrected by the replacement of the WDR domain with either the UBD_Doa1_ or NZF_Tab2_^31^. Consistently, we found here that both the UBD_Doa1_ and NZF_Tab2_ constructs significantly restored the WT pattern of intracellular structures labeled with mNG-Snc1 **(Fig. 2b, 2e)**. However, even though Snc2 is functionally and evolutionarily closely related to Snc1, we found that UBD_Doa1_ restored the mNG-Snc2 WT pattern, but the K63-restricted NZF_Tab2_ domain did not **(Extended data Fig. 6)**. For GFP-Sec22, β’COP-UBD_Doa1_ fully prevented vacuolar mislocalization while a partial rescue was conferred by β’COP-NZF_Tab2_ (**Extended data Fig. 1c-d).** Thus, Snc1 and Sec22 can use K63-linked polyUb chains for their trafficking, but Bet1, Gos1, and Snc2 appear to rely on COPI binding to some other ubiquitin linkage type.

Next, we examined the localization of mNG-tagged Bet1, Snc1, and Snc2 in a temperature-sensitive COPI mutant (*ret1-1)* grown at the permissive temperature and shifted to the non-permissive temperature of 37°C for 1 hr. The *ret-1* mutation is within α-COP and substantially inactivates all known COPI functions^5, 14, 36^. Bet1, Snc1, and Snc2 were mislocalized to tubular and ring-like structures in *ret1-1* at the non-permissive temperature **(Fig. 2a-f, Extended data Fig. 6)**. Interestingly, the mislocalization pattern seen for Bet1, Snc1, and Snc2 in β’COP Δ2-304 cells was comparable to *ret1-1* at the non-permissive temperature (**Fig1. b-c, Fig. 2a-c**). These data indicate that perturbations in the ability of β’-COP to bind ubiquitin in β’-COP Δ2-304 substantially disrupt COPI function with respect to Bet1, Snc1, and Snc2 localization. We previously showed that β’-COP Δ2-304 does not perturb Golgi to ER trafficking of cargoes bearing the KKXX or HDEL motifs^31^. Thus, it is the ability of the β’-COP N-terminal WDR domain to bind ubiquitin, not dilysine motifs, that is critical for SNARE localization.

β’-COP has been shown to bind K63 polyubiquitin (polyUb) chains but not K48 polyUb or monoubiquitin (monoUb)^31^. Since a general ubiquitin-binding domain rescued the localization for all 4 SNAREs, but not K63-specific ubiquitin-binding domain **(Fig. 2a-f)**, we reasoned that β’-COP might be able to bind other polyUb chains. To test this hypothesis, we assayed the ability of heterologously purified GST-tagged β’COP to bind K6-, K11-, K29-, K33-, and linear (M1)-linked polyUb chains. K63-polyUb was used as a positive control, and GST-only was used to determine background levels of ubiquitin-binding to GST (**Fig. 2g-h**). β’COP is capable of binding linear ubiquitin chains and more weakly to K6-, K11- and K29- polyUb chains (**Fig. 2g-h**).

### Fusion of a deubiquitinase domain to COPI leads to SNARE mislocalization

To analyze the functional significance of ubiquitination within the COPI-SNARE system, we designed constructs where a deubiquitinase domain, UL36 (DUB) from Herpes Simplex Virus 1^37^, was fused to either α-COP or β’COP. A catalytically dead version of UL36 (DUB*) wherein an active site Cys is mutated to Ser and thus cannot deubiquitinate substrates was engineered as a control. Strains expressing COPI-DUB constructs, irrespective of whether α-COP or β’COP was fused to DUB, were enlarged in size **(Fig. 3a, b)**. Additionally, we observed mislocalization of Bet1 and Gos1 in COPI-DUB constructs wherein mNG-tagged SNAREs were observed in enlarged punctate structures, elongated tube structures, or ring-like structures **(Fig. 3a, c)**. COPI-DUB* constructs did not display significant phenotypic changes. The fusion of a DUB domain to COPI phenocopies the mislocalization pattern for Bet1 and Gos1 in the COPI (*ret1-1*) mutant at nonpermissive temperatures, supporting the importance of ubiquitination in COPI mediated regulation of SNARE localization.

**Figure 3.**
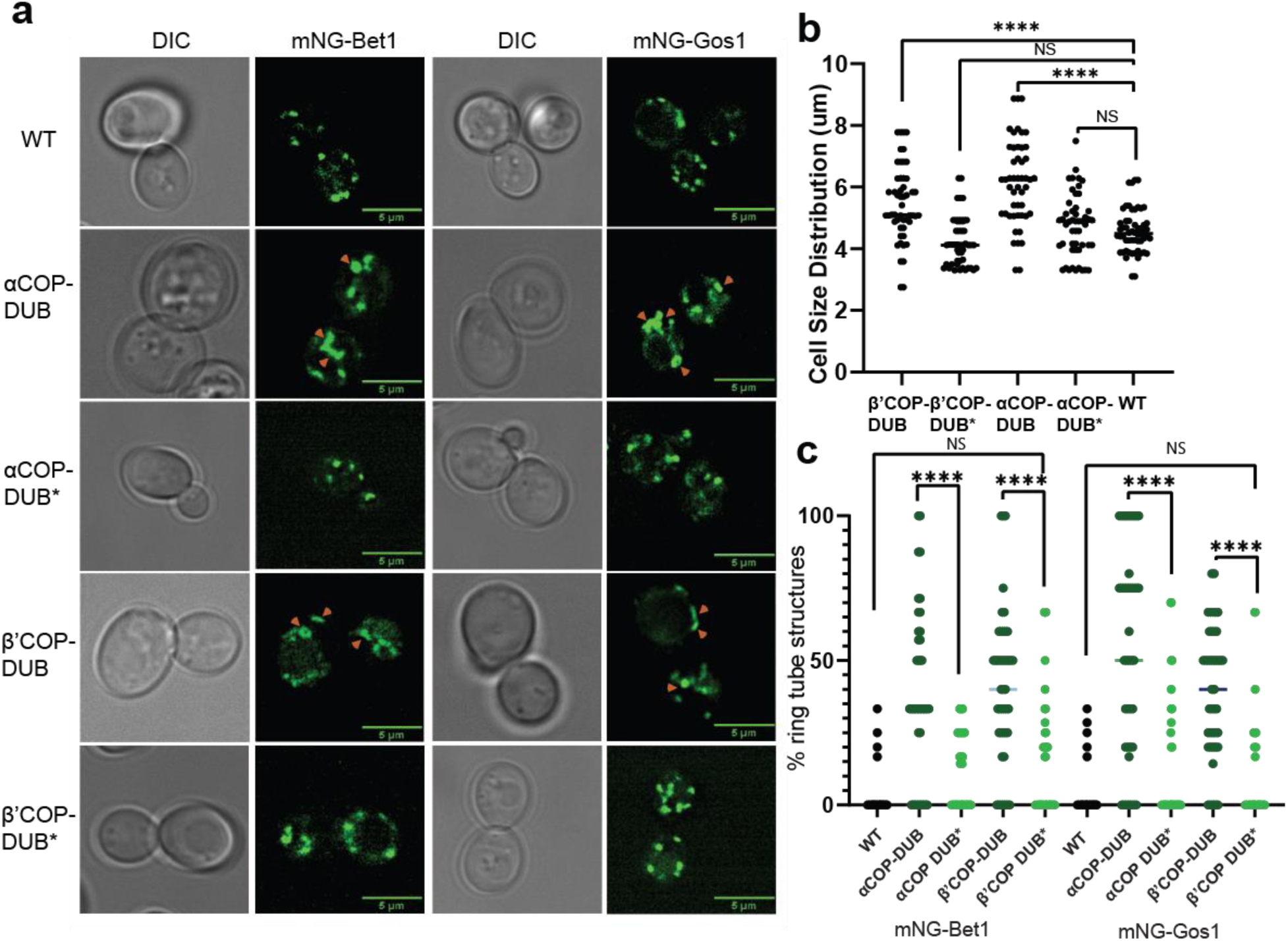
Deubiquitinase fusion to COPI subunits causes SNARE mislocalization. mNG-tagged Bet1 and Gos1 were imaged in cells in which a deubiquitinase domain (DUB) was fused to the C-terminus of α- and β’COP to generate αCOP-DUB and β’COP-DUB, respectively, along with catalytically dead controls αCOP-DUB* and β’COP-DUB*. **(a and b)** Cells carrying COPI-DUB fusion were larger in size compared to WT cells as well as catalytically dead controls. **(a and c)** Significant accumulation of Bet1 and Gos1 in the elongated tube- or ring-like or enlarged punctate structures (orange arrows) was observed in αCOP-DUB and β’COP-DUB backgrounds compared to corresponding DUB* control or WT cells. Statistical differences were determined using a one-way ANOVA on the means of the three biological replicates (***p<0.001).

### Ubiquitination is associated with Gos1, Ykt6, and Sed5 SNARE complexes

Global analyses of the budding yeast proteome have identified ubiquitinated lysines in Gos1, Snc1, and Snc2 but not Bet1^38^. We set out to test if ubiquitination could be detected by immunoprecipitating the SNAREs and probing for ubiquitin on immunoprecipitated samples and by detecting the pooled ubiquitin released off of immunoprecipitated samples following a deubiquitinase (DUB) treatment. We individually tagged Bet1, Gos1, and Snc1 with 6xHIS-TEV-3xFLAG at their C-termini by chromosomal integration of the tag constructs. Following FLAG immunoprecipitation, the samples were treated with mock buffer (no DUB) or deubiquitinases (DUB) (**Fig. 4a)** and probed with FLAG **(Fig. 4b)** or ubiquitin antibodies **(Fig. 4c)**. Art1, a ubiquitinated protein from *S. cerevisiae*, was used as a positive control, and untagged cells (Ctrl) were used as a negative control. The FLAG antibody recognizes a nonspecific band at approximately 20 kDa (**Fig. 4b, Ctrl Lane**) that unfortunately co-migrates with Bet1-FLAG and Snc1-FLAG as indicated by the increased band intensity at 20 kDa in those samples relative to the untagged control (Ctrl) sample. In addition, Bet1-FLAG exhibited a significant smear extending to greater than 40 kDa (**Fig. 4b**). However, this smeared pattern for Bet1-FLAG was not collapsed by DUB treatment, nor was this smear recognized by the anti-ubiquitin antibody. Moreover, the amount of monoUb released from Bet1-FLAG by DUB treatment was not significantly different from the control sample (**Fig. 4c-d**).

**Figure 4.**
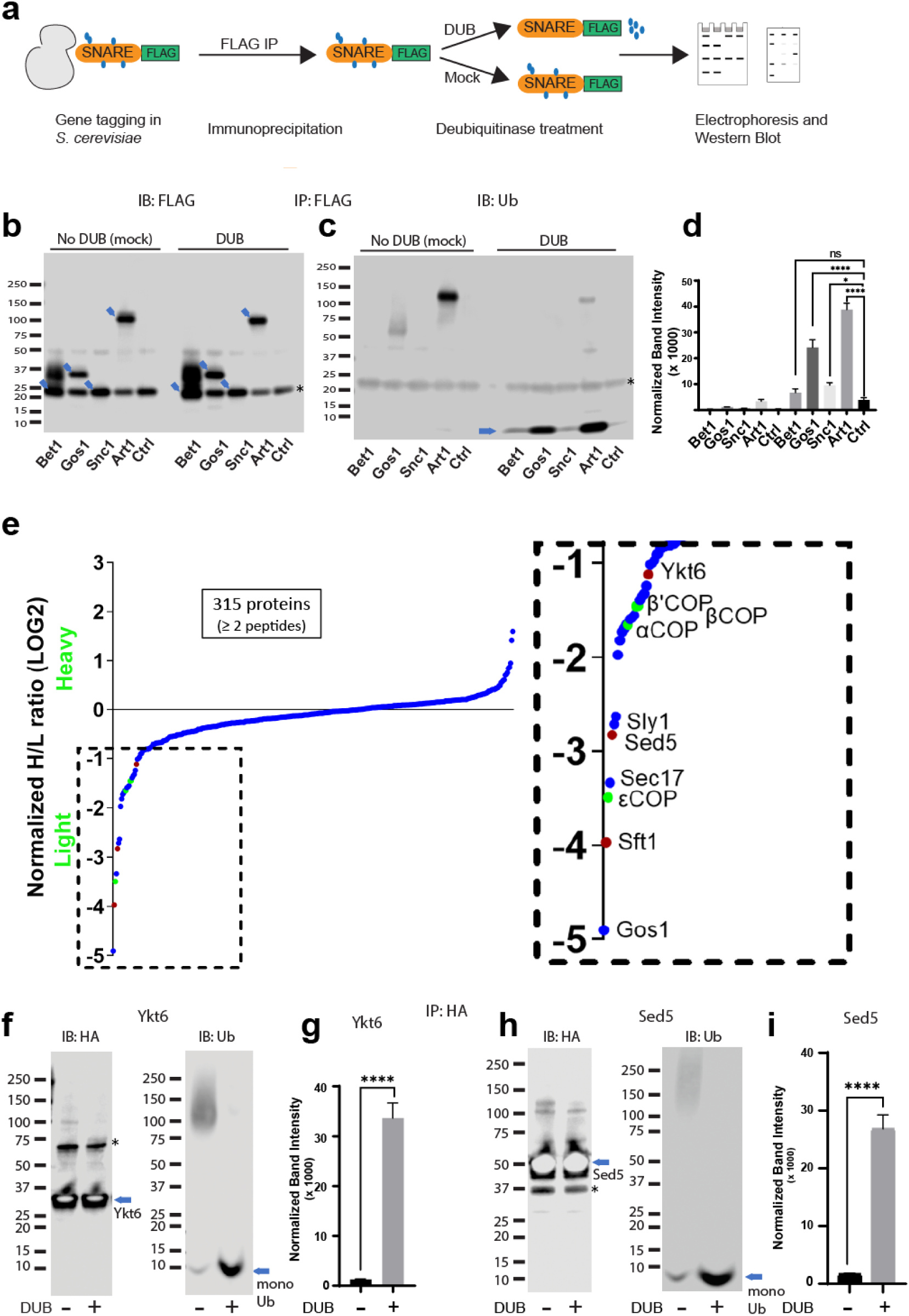
Multiple Golgi SNARE complexes are modified with ubiquitin. **(a)** Schematic of the experimental setup wherein SNAREs were individually tagged with FLAG and immunoprecipitated using anti-FLAG beads. Half the samples were mock-treated and the other half was treated with deubiquitinase (DUB). Western blots of samples are probed with FLAG **(b)** or ubiquitin antibody **(c)**. Blue arrows in (b) indicate the position of FLAG-tagged protein and the asterisk indicates the position of a background band. **(d)** Quantitation of the amount of monoubiquitin released from the samples by deubiquitinases. **(e)** SILAC mass spectrometric analysis of Gos1-FLAG pulldown samples indicates enrichment of SNAREs Sft1, Ykt6 and Sed5 (red dots) and COPI subunits (green dots) with Gos1. **(f-i)** Western Blot analysis showing HA-tagged Ykt6, and Sed5 probed for ubiquitination following HA-immunoprecipitation and deubiquitinase treatment. **(g,i)** Quantitation of monoubiquitin. Statistical differences were determined using a one-way ANOVA with multiple comparison test on three biological replicates (****p ≤ 0.0001, ***p<0.001, *p<0.05, Ns p > 0.05).

For Gos1-FLAG immunoprecipitations probed with anti-ubiquitin antibody, a smeared pattern was observed in the 50-80 kDa molecular weight region (**Fig. 4b**) when probed with anti-Ub, which collapsed, releasing a significant amount of monoUb following DUB treatment **(Fig 4c-d, Gos1 lanes)**. A similar smeared pattern is seen for Art1 in mock-treated samples around 75-130 kDa molecular weight region, which was converted to monoUb by DUB treatment **(Fig 4c-d, Art1 lanes)**. Although the smeared pattern for Snc1 was not apparent in these samples, DUB treatment released more monoUb than control samples **(Fig 4d, Snc1)**. We initially focused our attention on Gos1 because it appeared to be ubiquitinated and evidence for the importance of Snc1 ubiquitination has already been reported^31, 39^ **(Fig. 4c-d)**.

To identify other proteins specifically associated with Gos1 when purified under conditions that preserved ubiquitination, we employed a Stable Isotope Labeling by/with Amino acids in Cell culture (SILAC) mass spectrometry approach. A strain expressing Gos1-FLAG was grown in a light isotope medium and untagged control cells used to determine the nonspecific background proteins in the FLAG IP, were grown in a heavy isotope medium. Importantly, the samples were processed in the presence of DUB- inhibitors to preserve ubiquitination on Gos1 and other proteins in the samples. Gos1 is reported to form a functional t-SNARE complex with Ykt6 and Sed5 that mediates fusion with intra-Golgi retrograde vesicles bearing Sft1^40^. We observed significant enrichment of peptides from these partner SNAREs with Gos1-FLAG and known SNARE regulators like Sec17 and Sly1^41, 42^ **(Fig. 4e)**. Importantly, we also found several COPI subunit peptides that were enriched to comparable levels as Ykt6, Sft1, and Sed5 in the Gos1 pulldown samples **(Fig. 4e)**.

To probe the ubiquitination status of Gos1-binding SNARE partners, we individually tagged Ykt6 and Sed5 with 3xHA tag on the N-terminus (attempts at C-terminally tagging Ykt6 and Sed5 were unsuccessful potentially owing to structurally/functionally important modifications at the C-terminus, such as Ykt6 palmitoylation). HA-tagged Ykt6 and Sed5 were immunoprecipitated using anti-HA and probed for their ubiquitination status. A smeared pattern associated with ubiquitination was observed for both Ykt6 (**Fig. 4f**) and Sed5 (**Fig. 4h**) in mock-treated samples, which was collapsed by DUB treatment to monoUb **(Fig. 4f-i)**. These data support previously published high-throughput results indicating that Gos1, Ykt6, and Sed5 are ubiquitinated^38^. The differences in the size distributions of polyUb smear in each SNARE immunoprecipitate suggest that this assay is primarily detecting direct modification of Gos1, Ykt6, and Sed5 as opposed to the aggregate polyUb associated with the entire SNARE complex.

### Non-degradative ubiquitination is associated with Gos1, COPI, and Glo3 complexes

We observed significant enrichment of COPI subunits in the Gos1 pulldown samples analyzed with SILAC mass spectrometry **(Fig. 4e)**. Therefore, we probed the ubiquitination status of FLAG-tagged COPI (α- and β’COP subunits) and Glo3, as this ArfGAP is reported to bind COPI and SNAREs^19^. The FLAG IPs probed with anti-ubiquitin antibody show a substantial amount of monoUb released from COPI and Glo3 immunoprecipitates following the DUB treatment **(Extended Data Fig. 7a-e)**. K48- linked polyUb chains are known to target proteins for proteasomal degradation. To address whether Gos1, COPI, and Glo3 complexes are modified with K48-linked polyUb, we treated the samples with a K48- specific DUB. No significant change in the smeared electrophoretic pattern or the release of monoUb in the samples was observed with or without K48-specific DUB treatment suggesting that the ubiquitination associated with COPI and Glo3 is not a degradation signal **(Extended Data Fig. 7a-e)**.

We also probed Gos1, Ykt6, Sed5, COPI, and Glo3 FLAG-immunoprecipitated samples with K63 specific deubiquitinase (K63-DUB^43^), linear-ubiquitin specific deubiquitinase (M1-DUB^44^) or a general deubiquitinase (DUB^45^) as a control **(Extended Data Fig. 8)**. No significant release of ubiquitin was observed following K63-or M1-DUB treatment compared to the untagged control **(Extended Data Fig. 8)**. A detectable amount of ubiquitin was released from Gos1 following K63-DUB treatment, but the signals were not significantly above the background levels **(Extended Data Fig. 8)**. A significant level of released ubiquitin was detected for these samples when treated with the general deubiquitinase. The lack of K63 linkages on these components is also consistent with live-cell imaging data **(Fig. 2a-f)**, showing that β’-COP with a K63-specific binding domain failed to support the trafficking of Bet1, Gos1, and Snc2. Thus, the ubiquitination associated with Gos1, COPI, and Glo3 complexes appears to be non-degradative (non-K48 or non-K63) in nature and may modulate protein interactions in the COPI-dependent retrieval of SNAREs within the Golgi.

### Ubiquitination stabilizes Golgi SNARE-COPI complexes

To explore the possibility that ubiquitination is an important regulator of protein-protein interactions in the COPI-SNARE system, we used comparative pulldown studies using FLAG-tagged SNAREs under conditions that preserved endogenous ubiquitination (w Ub) or catalyzed removal of ubiquitin (w/o Ub) (**Fig. 5a**). An equal amount of Gos1-FLAG was pulled down in both w Ub and w/o-Ub conditions **(Fig. 5c)**. Probing samples with a ubiquitin antibody showed a ubiquitin smear associated with Gos1 immunoprecipitated using ‘w Ub’ conditions, most of which was stripped off under ‘w/o-Ub’ conditions **(Fig. 5b)**. We next probed these samples with COPI and Arf antibodies. Significant enrichment of COPI subunits and Arf was observed with Gos1 when ubiquitination was preserved, compared to ‘w/o-ub’ conditions **(Fig. 5 e-g).** The ubiquitination, thus, appears to play a role in the assembly and/or stability of COPI coatomer complex with Gos1.

**Figure 5.**
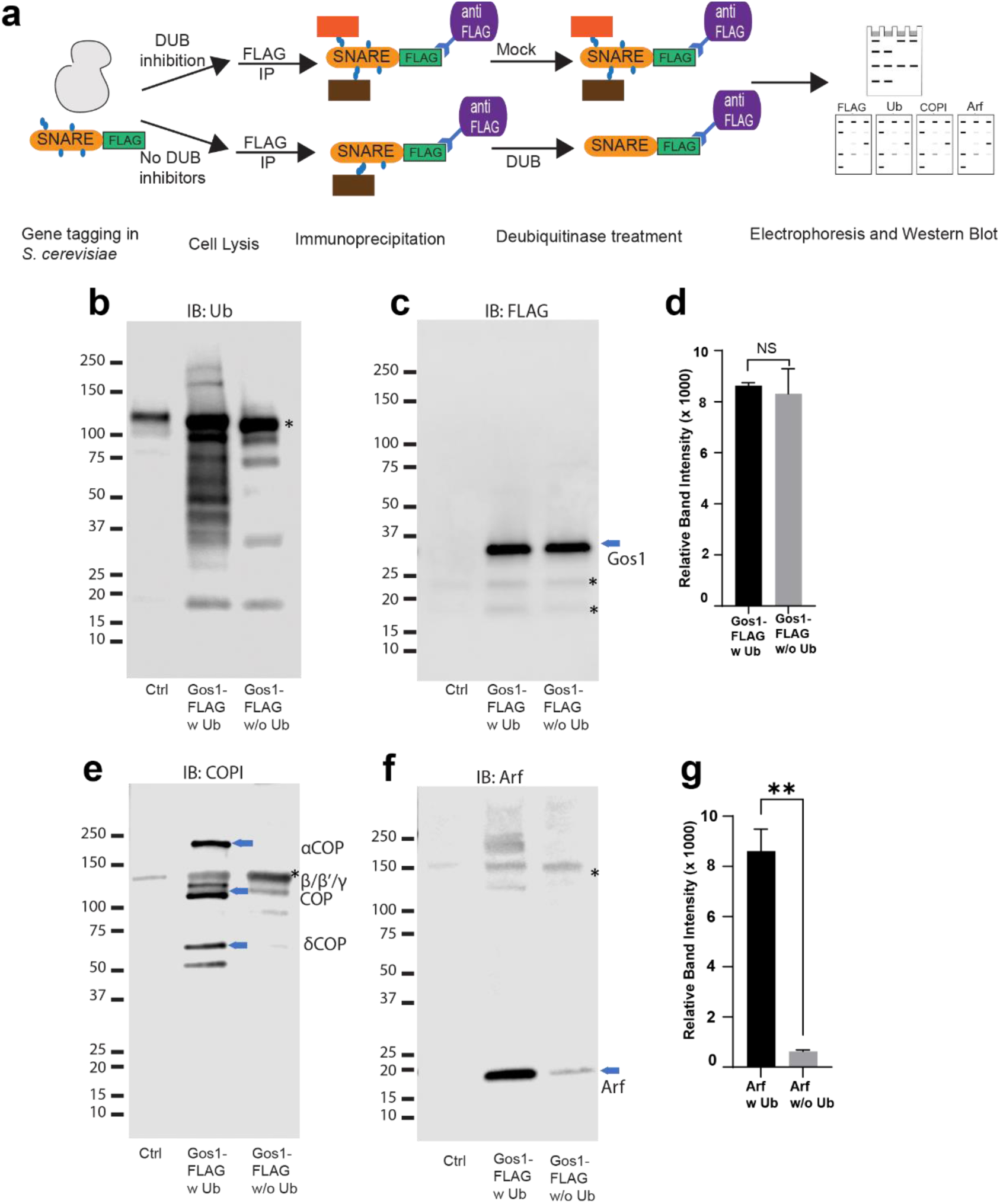
Ubiquitin modification stabilizes a priming complex between COPI, Arf and SNAREs. **(a)** Schematic of the experimental setup wherein FLAG-tagged SNAREs are divided into two equal portions, and one is processed under ‘w Ub’ conditions (DUB inhibitors used during the lysis step and no deubiquitinases (mock) treatment) and the other portion is processed under ‘w/o Ub’ condition (no deubiquitinase inhibitors used during lysis and immunoprecipitated samples are treated with deubiquitinases). **(b-i)** Western Blot data showing comparative pulldowns of Gos1-FLAG **(b-e)** and Bet1-FLAG **(f-i)** processed under ‘ubiquitin-preserved’ (w Ub) and ‘no-ubiquitin’ (w/o Ub) condition, and probed for Ub **(b,f)**, FLAG **(c,g)**, COPI **(d,h)** and Arf **(e,i)**. Untagged cells processed under ‘w UB’ condition to determine background binding were used as a control (Cntr) and abundant background bands are marked with an asterisk. Quantitation of **(j-k)** Gos1-FLAG and Arf, and **(l, m)** Bet1-FLAG and Arfin the pulldown samples. Band intensities are measured using ImageJ. Quantitation was done on three biological replicates using a t-test (****p ≤ 0.0001, ***p<0.001, *p<0.05, Ns p > 0.05).

Our assays and previous reports indicate that Gos1 is ubiquitinated^38^, but ubiquitination has not been detected on Bet1 (**Fig. 4b-d**). Loss of the ubiquitin-binding domain of COPI in β’-COP Δ2-304 led to mislocalization of both Bet1 and Gos1 to the elongated tube- and ring-like structures (**Fig.1b, Fig. 2a,c**). Therefore, we tested whether ubiquitination affects the interaction of Bet1 with COPI coat complex components. The smeared pattern associated with Bet1 in the blot probed with anti-FLAG antibody is similar under ‘w Ub’ and w/o Ub’ conditions (Extended Data **Fig 9b-c**). Nonetheless, we see the enrichment of COPI subunits with Bet1 under ‘w Ub’ conditions compared to ‘w/o Ub’ (Extended Data **Fig. 9d**). Similarly, Arf is significantly enriched with Bet1 when ubiquitin was present on the complexes (Extended Data **Fig. 9e-f**). Control experiments indicated that the presence of DUB inhibitors during cell lysis was most critical to preserve the SNARE-COPI complex **(Fig.5a, Extended Data Fig 9g,h)**. Therefore, the role of ubiquitination in the assembly and stability of COPI coatomer complex with Bet1 appears to function independently of the Bet1 ubiquitination status. Altogether, the data reveal ubiquitin-mediated stabilization of COPI-Golgi SNARE complexes.

### Glo3 is not enriched in ubiquitin-stabilized SNARE-COPI-Arf complexes and Gos1 localization is unaffected in *glo3Δ* cells

Glo3 is proposed to be part of a SNARE-Arf-COPI priming complex but we failed to detect any Glo3 peptides in the Gos1 immunoprecipitates by mass spectrometry (**Fig. 4e**). To further test whether ArfGAP Glo3 is present in the ubiquitin-stabilized SNARE-COPI-Arf complex, we performed Gos1-FLAG pulldowns under w Ub and w/o-Ub conditions in cells expressing Glo3 C-terminally tagged with GST. We detected a small amount of Glo3-GST co-immunoprecipitating with FLAG-Gos1, but no significant difference was observed in the presence or absence of Ub **(Fig. 6a,c)**. In contrast, association of Arf with Gos1 was significantly enriched using w Ub conditions compared to w/o-Ub conditions (**Fig. 6a,d**). Cell lysate controls probed for GST in cells expressing only Gos1-FLAG or both FLAG-Gos1 and Glo3-GST confirmed the identity of the Glo3-GST band (**Fig. 6b**).

**Figure 6.**
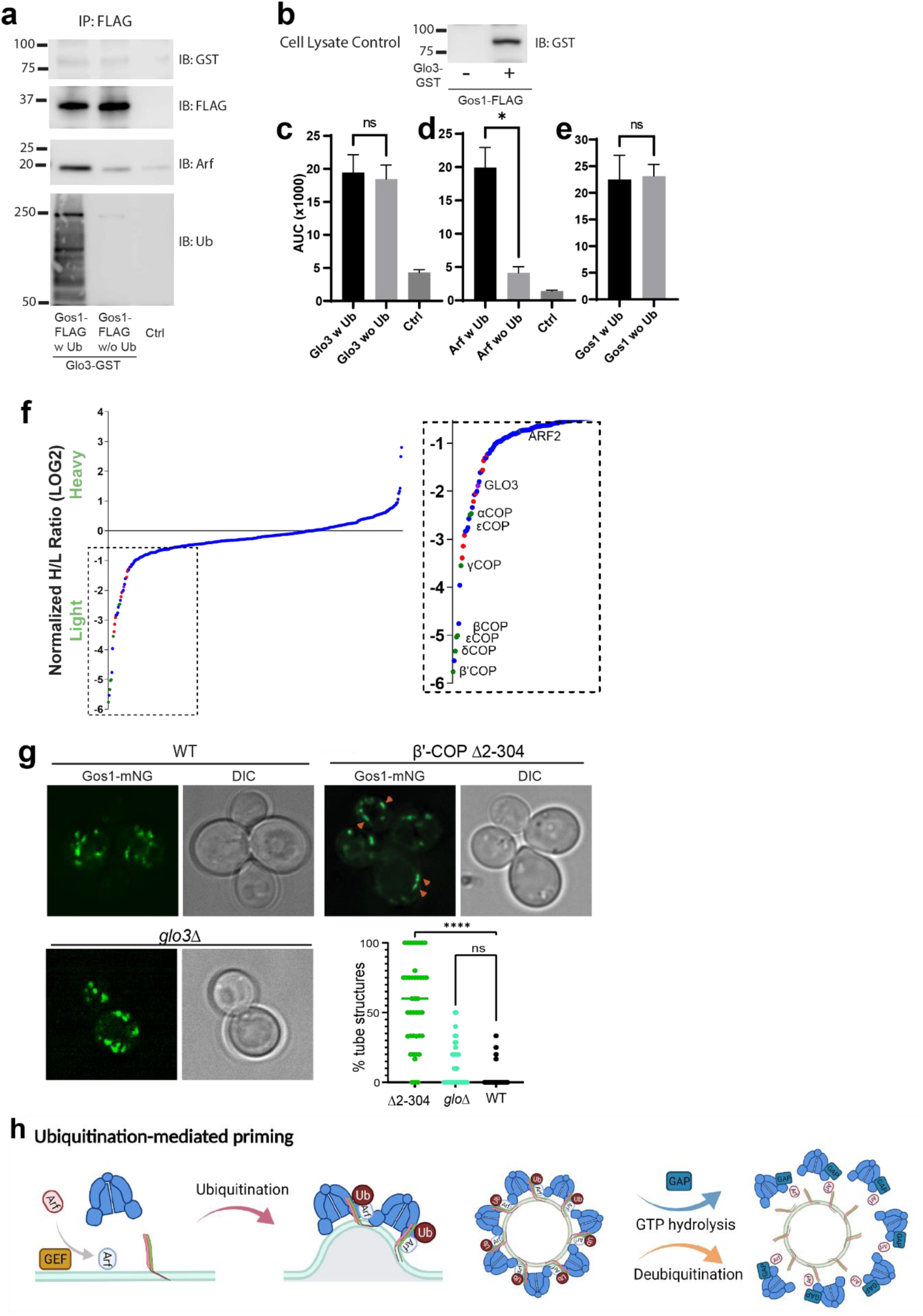
ArfGAP is not enriched in ubiquitin-stabilized SNARE-Coat complexes and is not required for Gos1 localization. (a-e) Western Blot data showing comparative pulldowns of Gos1-FLAG from cells expressing FLAG-tagged Gos1 and GST-tagged Glo3. Samples were processed under ‘ubiquitin-preserved’ (w Ub) and ‘no-ubiquitin’ (w/o Ub) condition, and probed for Glo3 (anti-GST), Gos1 (anti-FLAG), Arf, and Ub. Untagged cells processed under ‘w UB’ condition to determine background binding were used as a control (Ctrl). Cell lysates from cells expressing only FLAG-tagged Gos1 or both FLAG-tagged Gos1 and GST-tagged Glo3 probed with anti-GST antibody are included as controls to ensure expression of GST-tagged Glo3. Quantitation of **(c)** Glo3-GST, **(d)** Arf and **(e)** Gos1-FLAG samples. Band intensities are measured using ImageJ. Quantitation was done on three biological replicates using a t-test (*p<0.05, Ns p > 0.05). **(f)** SILAC mass spectrometric analysis of αCOP-FLAG pulldown samples indicate enrichment of other COPI subunits (green dots), ArfGAP Glo3 (purple dots) and diLys COPI cargo (red dots) but no SNAREs. **(g)** Live cell imaging of mNG-Gos1 in WT, β’COP Δ2-304 and *glo3Δ* cells. Quantification of % tube structures for each strain was from 3 biological replicates with 60 or more cells analyzed for each sample. Statistical differences were determined using a one-way ANOVA on the means of the three biological replicates (***p<0.001). Scale bar in represents 5 µm. **(h)** Model showing ubiquitination-mediated priming of a SNARE-Arf-COPI complex. Glo3 is recruited at later stages after vesicle budding leading to hydrolysis of Arf-GTP and disassociation of COPI complex.

To determine whether we can detect Glo3 interaction with COPI, we performed SILAC coimmunoprecipitation analysis using C-terminally FLAG-tagged α-COP under ubiquitin preserved conditions. As expected, Glo3 was enriched with COPI, indicating eventual recruitment of Glo3 onto the COPI coat **(Fig.6f)**. The COPI sample was also enriched for several ER residence membrane proteins bearing C-terminal dilysine motifs (Fig. 6f, red datapoints). However, no SNAREs were coimmunoprecipitated with COPI, indicating that the COPI complexes recovered here were significantly different from SNARE-COPI complexes that contained Gos1.

In addition, we analyzed mNG-Gos1 localization in *glo3Δ* cells with WT and β’COP Δ2-304 cells as control. Compared to WT cells, no significant changes in the size or distribution of mNG-Gos1 punctae were observed in *glo3Δ* (Fig. 6g). As reported earlier, Gos1 is mislocalized to elongated tube-like structures in β’COPΔ2-304 cells. Together, these data suggest that Glo3 can weakly bind Gos1 but unlike COPI, this interaction is not stabilized by the presence of Ub. In addition, Glo3 does not appear to be required for COPI-dependent Gos1 localization.

Altogether, these results indicate that ubiquitin plays critical role in stabilizing a complex between SNAREs, COPI and Arf that is important for COPI function in retrieving a subset of SNAREs. Glo3 appears to be mostly excluded from the ubiquitin-stabilized Gos1-Arf-COPI complex

## Discussion

We previously discovered that COPI binds specifically to polyUb chains and that this interaction is crucial for recycling Snc1, an exocytic v-SNARE, back to the TGN. In this study, we broadly probed the role of COPI-ubiquitin interactions on the localization of 16 additional budding yeast SNAREs to determine whether ubiquitination of coat components is a general mechanism for SNARE sorting. While localization of most mNG-tagged SNAREs was unaffected by deletion of the ubiquitin-binding N-terminal WDR domain of β’-COP, we found a significant change in the localization pattern for Bet1, Gos1, Snc1, Snc2 and partially for Bos1 and Sec22 in β’-COP Δ2-304 (**Fig. 1b-c, Fig. 2 a-f, Extended Data Fig. 1**). Normal SNARE localization is restored by the replacement of β’ COP N-terminal WDR domain (β’ COP-UBD) with an unrelated ubiquitin-binding domain (**Fig. 2, Extended Data Fig. 6**). Moreover, we found non-degradative (non-K48) ubiquitination associated with multiple Golgi SNAREs (Gos1, Ykt6, Sed5) and the COPI machinery (**Fig. 4b-d, f-i, Extended Data Fig.7, 8**), and that these ubiquitin modifications were essential for the stabilization of COPI, Arf, and Golgi SNARE complexes (**Fig. 5**). For Gos1, the ubiquitin-stabilized SNARE-COPI-Arf complex lacks the ArfGAP Glo3 **(Fig.6)**. These studies highlight the important role of ubiquitination in COPI-mediated trafficking, specifically in the regulation of Golgi SNARE localization.

The type of ubiquitin linkages required for intra-Golgi SNARE interaction with COPI appears to be different from the ubiquitin linkages required to sort Snc1. β’-COP binds preferentially to K63-linked polyUb chains and does not bind monoUb, diUb or K48-linked polyUb chains^31^. Replacement of the N-terminal WDR domain of β’-COP with the NZF domain from Tab2, which binds specifically to k63- linked polyUb, substantially restores Snc1 trafficking to the plasma membrane^31^ **(Fig. 2b)**. Surprisingly, the β’-COP-Tab2_NZF_ fusion fails to support the trafficking of Bet1 or Gos1 and only partially supports the trafficking of Sec22 (**Fig. 2a-f, Extended Data Fig. 1**). We further explored the binding specificity of β’- COP and found that it is also capable of binding linear polyUb chains and to K6-, K11- and K29- polyUb chains (**Fig. 2g-h**). Moreover, the polyUb chains detected in the SNARE or COPI pulldowns are resistant to M1-, K63- or K48-specific DUBs (**Extended Data Fig. 8**). Thus, the ubiquitin modifications present are unlikely to be targeting COPI to the proteasome or the SNAREs to the vacuole for degradation. Our data provide compelling evidence that the non-proteolytic ubiquitin code regulates the COPI-dependent trafficking patterns for Golgi SNAREs.

PolyUb chains on SNAREs could form a sorting signal that COPI recognizes in order to recycle them from downstream compartments, as previously proposed for Snc1^31^. However, several observations in the current study suggest broader roles of ubiquitination in regulating SNARE trafficking. For example, the medial Golgi localization of Bet1 relies on COPI’s ability to bind ubiquitin, but we could not detect ubiquitinated forms of this SNARE **(Fig. 4b-d)**. However, ubiquitination was associated with other SNAREs, including Gos1, Ykt6, and Sed5, multiple COPI subunits, and ArfGAP Glo3 **(Fig. 4b-d, f-i, Extended Data Fig. 7, 8)**. It is possible that Bet1 associates with another cargo protein that is ubiquitinated, and the ubiquitin serves as the COPI-dependent sorting signal for both proteins. It is also possible that ubiquitination induces conformational changes in COPI driven by β’-COP interaction with ubiquitin attached to itself or to other COPI subunits. Such a COPI conformational change could produce a high-affinity binding site for Bet1. The role of ubiquitin in mediating the stability of the COPI-SNARE complex is further supported by the observation that Bet1 and Gos1 are mislocalized when ubiquitin is stripped from COPI-SNARE system by fusing a deubiquitinase domain to COPI components **(Fig 3)**.

We were surprised to find that COPI was co-enriched with Gos1-FLAG in the SILAC-based mass spectrometry data (**Fig. 4e**) because cargo-coat interactions are typically low affinity. Arf1 was also present in this dataset, although not as highly enriched as the COPI subunits. We considered the possibility that the conditions used to pulldown Gos1-FLAG that preserve ubiquitination may have stabilized the COPI-Gos1 interaction. Indeed, performing these Gos1-FLAG pulldowns in the presence of active DUBs to remove ubiquitin dramatically reduces the amount of COPI and Arf recovered with Gos1- FLAG relative to samples prepared with DUB inhibitors present (**Fig. 5e-g**). The interaction between Gos1 and COP1/Arf is nearly undetectable if ubiquitination of the components is not preserved. Bet1 interaction with Arf/COPI is also enhanced substantially under conditions that preserve ubiquitination (Extended Data **Fig.9d-f**). Therefore, ubiquitination appears to regulate the assembly and/or stability of the COPI-cargo complex independent of the ubiquitination status of cargo. Not all ubiquitinated SNAREs relied on COPI-ubiquitin interaction for their sorting, For example, Sed5 is ubiquitinated but its localization not affected by the alterations in the ability of COPI to recognize and bind ubiquitin (**Fig. 4 h-i, Extended Data Fig. 1, Sed5**), and Sed5 appears to be independent of COPI for its Golgi localization^24^. A subset of Golgi SNAREs is dependent on the ability of COPI to bind ubiquitin (**Extended Data Fig. 1**), and it is likely that other ubiquitin-independent interactions contribute to cargo selection.

Our data provide an exciting window into understanding the molecular details of organelle homeostasis in cells, particularly Golgi biology. SNARE trafficking patterns must play a critical role in establishing the organization and function of the Golgi complex. Bet1 is a v-SNARE that forms a fusogenic SNARE complex with the early Golgi syntaxin Sed5, R-SNARE Sec22, and Bos1^15, 46, 47^. Sed5 and Sec22 are ubiquitinated and could possibly facilitate retrieval of Bet1 in COPI vesicles. However, the trafficking patterns for these SNAREs are different. Bet1 does not recycle back to the ER as one would expect if Bet1 was serving as the v-SNARE in COPII vesicles. In contrast, Sed5, Sec22 and Bos1 do recycle through the ER, and therefore, it is possible that this trimeric complex is the active fusogenic SNARE in COPII vesicles budding from the ER ^33, 40, 48^. A key event in Golgi biogenesis may be the fusion of ER-derived COPII vesicles bearing Sed5-Sec22-Bos1 with Golgi-derived COPI vesicles bearing the v-SNARE Bet1 and also carrying early Golgi enzymes.

One of the long-standing questions about COPI-mediated vesicular trafficking has been the essential roles of α and β’COP WDR domains. α and β’COP WDR domains are essential for the sorting of dilysine motif COPI cargoes, but cells are viable when all dilysine sites mutated^13^. These studies indicated possible additional roles of COPI WDR domains in cells. Our studies address this critical question by showing the essential role of β’COP WDR in binding ubiquitin and mediating localization of ubiquitinated cargoes.

Another key element of vesicle-mediated trafficking is the ability of the coat to bind cargo during vesicle formation, followed by dissociation after the vesicle forms. SNAREs are thought to prime coat assembly through interactions with Arf, ArfGAPs, and COPI as a mechanism to ensure vesicles form with an adequate load of v-SNAREs^19, 49^. The ArfGAP Glo3 contains a BoCCS motif that mediates binding to both COPI and to several different SNAREs, suggesting that Glo3 is a key determinant of the priming complex^50^. However, it is unclear how Glo3 could facilitate coat assembly when its enzymatic function is to inactivate Arf. We have identified a ubiquitin-stabilized complex between Gos1, Arf and COPI that lacked endogenous Glo3 (**Fig. 4**). We were able to detect a tagged form of Glo3 in Gos1 immunoprecipitates that lack ubiquitin; however, preserving ubiquitin in these Gos1 pulldowns had no influence on Glo3 recovery even though substantially more Arf and COPI were recovered. The presence of ubiquitin modifications on COPI and Glo3 does not prevent their interaction because we observed enrichment of Glo3 in COPI pulldowns under the same ubiquitin-preserved conditions. Therefore, we suggest that the SNARE/Arf-GTP/COPI priming complex is stabilized by ubiquitination of the components and is devoid of ArfGAP. Arf-GDP and COPI likely dissociate rapidly from the complex as the ArfGAP binds **(Fig.6h)**.

Ubiquitin-dependent enrichment of Arf and COPI with SNAREs suggests that cycles of ubiquitination and deubiquitination could control the switch from Arf-GTP/SNARE-mediated assembly of COPI during budding and ArfGAP-mediated disassembly and uncoating of vesicles prior to fusion. Ubiquitination-deubiquitination cycles for key components within the COPI-SNARE system thus may alter the coatomer assembly-disassembly dynamics regulating COPI function.

## Methods

### Reagents

ANTI-FLAG M2 Magnetic Beads (M8823), EZview™ Red ANTI-FLAG® M2 Affinity Gel (F2426 ), 3xFLAG Peptide (F4799), N-Ethylmaleimide (E3876), Iodoacetamide (GERPN6302), 1,10-Phenanthroline (131377), N-Ethylmaleimide (E3876), deubiquitinase inhibitor PR-619 (SML0430), protease inhibitor tablets (04693159001), phosphatase inhibitors tablets (PHOSS-RO) were purchased from MilliporeSigma (St Louis, MO). Coomassie Brilliant Blue R-250 Dye (20278), and FM4-64 dye (T-3166) were purchased from ThermoFisher Scientific (San Jose, CA). ECL Prime Western Blotting Chemiluminescent Substrate (34580), Pierce™ Anti-HA Agarose (26181) were purchased from Thermo Scientific (Rockford, IL). Deubiquitinases (DUBs) Usp2 (E-504), MINDY2 (E-620), MINDY3 (E-621), OTULIN (E-558), AMSH (E-548B), K6-ubiquitin trimer (Ub3) chains (UC-20-025), K11-Ub3 chains (UC-50-025), K29-Ub3 chains (UC-85-025), and K33-Ub3 (UC-105-025) were from BostonBiochem- R&D Systems, Inc. (MN, USA)

### Antibodies

ANTI-FLAG® antibody produced in mouse (clone M2, F3165, 1:3500) and Anti-HA antibody produced in rabbit (H6908, 1:1000) were purchased from MilliporeSigma (St. Louis, MO). VU101: Anti-ubiquitin Antibody (VU-0101, 1:1000) was purchased from LifeSensors (PA, USA). Anti-mouse HRP conjugate (W4021, 1:10,000) and Anti-Rabbit HRP Conjugate (W4011, 1:10,000) were purchased from Promega (Madison, WI). Anti-COPI antibody was a gift from Charles Barlowe (Dartmouth Univ, 1:3000). Anti-Arf antibody (1:3000) used was reported previously^51^. Anti-GST antibody (1:1000) was purchased from Vanderbilt Antibody Product Store (VAPR, Nashville, TN).

### Strains and plasmids

Standard media and techniques for growing and transforming yeast were used. Epitope tagging of yeast genes was performed using a PCR toolbox ^52, 53^. The list of yeast strains used in this study are included as a table file (Supplementary file 1). Plasmid constructions were performed using standard molecular manipulation. Mutations were introduced using Gibson Assembly Master Mix. The list of plasmids used in this study is included as a table file (Supplementary file 2).

### Imaging and image analysis

To visualize mNeonGreen- or mScarlet-tagged proteins, cells were grown to early-to-mid-logarithmic phase, harvested, and resuspended in imaging buffer (10 mM Na_2_PHO4, 156 mM NaCl, 2 mM KH_2_PO4, and 2% glucose). Cells were then mounted on glass slides and observed immediately at room temperature. Images were acquired using a DeltaVision Elite Imaging system equipped with a 63× objective lens followed by deconvolution using SoftWoRx software (GE Healthcare Life Science). Overlay images were created using the merge channels function of ImageJ software (National Institutes of Health). To quantify SNAREs colocalization, a Pearson’s Correlation Coefficient (PCC) for the two markers in each cell (n =3, over 20 cells each) was calculated using the ImageJ plugin Just Another Colocalization Plugin with Costes Automatic Thresholding ^54^.

Identification and quantitation of fluorescence-based morphological patterns were performed as below: the punctate pattern indicates small, dotted structures, typically around 0.2 – 0.3 µm, the ring-like structures indicate larger, roughly donut-shaped structures approximately 2-4 times larger than the ‘punctate’ structures, and elongated tube-like structures indicate tube-like structures, approximately 2-5 times in length along the plane compared to the ‘punctate’ pattern. Each fluorescent structure in the cell was categorized as puncta, ring or tubule and the number of tubules + ring divided by total fluorescent structures was used to quantify the % tubular and ring structures. Measurements were done in minimum of 50 cells (n ≥50) for 3 biological replicates. Fluorescence pattern identification and quantitation were repeated in a blinded fashion and/or by a second observer to avoid bias.

### Synthesis of K63 and linear ubiquitin chains

To synthesize K63 linked Ub chains, 2 mM Ub, 300 nM E1, 3 μM UBE2N/UBE2V2 were mixed in the reaction buffer (50 mM Tris-HCl pH 7.5, 50 mM NaCl, 10 mM MgCl_2_, 20 mM ATP, and 2 mM DTT) overnight at 37°C. Reactions were quenched by lowering the pH to 4.5 with addition of 5 M ammonium acetate pH 4.4. K63 tri-Ub were isolated and further purified using size exclusion chromatography (Hiload 26/600 Superdex 75 pg, GE Healthcare) in gel filtration buffer (50 mM Tris-HCl pH 7.5, 150 mM NaCl, 1 mM DTT). Purified chains were buffer exchanged into H_2_O and lyophilized. Recombinant M1-linked tri-Ub-FLAG-6XHis was expressed and purified as previously described^55^. Briefly, *E. coli* C41(DE3) cells at OD_600_ of 0.6 were induced with 1 mM IPTG, lysed by sonication in ice-cold Tris buffer (50 mM Tris pH 8.0, 150 mM NaCl, 10 mM imidazole, 2 mM βME, complete protease inhibitors (Roche, Basel, Switzerland), 1 μg/ml DNase, 1 μg/ml lysozyme, and 1 mM PMSF), and clarified by centrifugation (50,000 x *g* for 30 min at 4°C) and filtration (0.45 μM filter). M1 tri-Ub was purified to homogeneity by Ni^2+^-NTA affinity column (Thermo Scientific, Rockford, IL) chromatophraphy, HiPrep Q FF anion exchange column (GE Healthcare Life Sciences, Marlborough, MA) chromatography, and HiLoad Superdex size-exclusion column (GE Healthcare Life Sciences, Marlborough, MA) chromatography.

### Construction FLAG-, HA- and GST-tagged constructs

Multiple strains of *Saccharomyces cerevisiae* were generated in a manner where one of the components is tagged with an epitope tag. Bet1, Gos1, Snc1 were C-terminally tagged with 6xHis-TEV-3xFLAG by integration of a PCR product amplified from pJAM617 into the *BET1, SNC1* and *SNC2* locus respectively ^52^. Due to the low recombination rate, a *GOS1* PCR product with longer 5’ and 3’ regions of homology (over 200bp) was generated from pJAM617 and gene synthesized DNA fragments and integrated into the *GOS1* locus (two-step PCR and integration method). Properly integrated clones were confirmed by genotyping PCR as well as by immunoblot using anti-FLAG antibody. Similarly, COP1, Sec27, ArfGAP Glo3 were C-terminally tagged with 6xHis-TEV-3xFLAG using 2-step PCR and integration method. Additionally, Glo3 was C-terminally tagged with GST (using pFA6a-GST-HisMX6 as a template) in WT *S. cerevisiae* as well as in cells harboring 6xHis-TEV-3xFLAG-tagged Gos1. Efforts to C-terminally tag Ykt6 and Sed5 were unsuccessful; consequently, Ykt6 and Sed5 were N-terminally tagged with 6xHA tag by integration of a PCR product amplified from pYM-N20 cassette (Euroscarf #P30294).

### Purification of FLAG-tagged or HA-tagged proteins

Affinity isolation of FLAG-tagged or HA-tagged proteins was performed with anti-FLAG magnetic beads or Anti-HA Agarose, respectively. 800 OD_600_ of untagged wild-type cells (BY4742) and cells with C- or N-terminally tagged protein of interest were grown in YPD and harvested by centrifugation when the OD_600_ reached ∼0.8. After washing with cold water, the pellets were resuspended in 3 mL lysis buffer (100 mM Tris pH 7.4, 150 mM NaCl, 5 mM EDTA, 5 mM EGTA, 10% glycerol, 1% Triton X-100, 100μM PR619, 5 mM 1,10-Phenanthroline, 50 mM N-Ethylmaleimide, phosphatase inhibitors and complete protease inhibitor tablet). Cells were broken using a Disruptor Genie (Scientific Industries) at 4°C for 10 min at 3000 setting with 0.5 mm diameter glass beads. The lysates were centrifuged at 13,000 rpm for 15 min at 4°C and the supernatant was incubated with 50µL FLAG or HA beads overnight at 4°C. The next morning the beads were washed 3x with washing buffer (100 mM Tris pH 7.4, 150 mM NaCl, 5 mM EDTA, 1% NP40, 0.5% Triton X-100) and eluted in SDS running buffer.

Heterologous expression and purification of GST-β’COP and ubiquitin binding assays were performed as reported previously^31^. Briefly, 0.5 mM of GST and GST tagged β’COP (604) proteins immobilized glutathione beads were incubated 250 nM ubiquitin trimer (Ub3) for corresponding linkages, washed 3x and eluted using reduced glutathione.

### DUB treatments

The DUB treatments were performed as described ^56^. Briefly, the beads with target proteins were equally split into two parts. One part was subjected to mock treatment as a control, and the other part was incubated with deubiquitinases in the following reaction: 5 μl of 10xDUB reaction buffer (1M Tris pH 7.4, 1.5 M NaCl, 10 mM DTT), 0.5 μl of deubiquitinase enzyme, and water for a 50 μl reaction volume. The samples were incubated at 37°C for 45 minutes and reactions were stopped with 2x Laemmli sample buffer by incubating at 95°C for 5 minutes. Supernatants were collected and used for electrophoresis followed by Western transfer. Deubiquitinases were used following manufacturer recommended concentrations as follows: DUB: the general deubiquitinase Ups2 (1-5 nM); K48-DUB: K48 linkage specific deubiquitinase MINDY2 and MINDY3 (10-30 nM); K63-DUB: K63 linkage specific deubiquitinase AMSH (100-500 nM); M1-DUB: OTULIN (0.05­ 1 μM). Data were generated from independent experiments from three biological replicates and quantified as described later.

For comparative pulldown samples processed under conditions that preserved endogenous ubiquitination (w Ub) or catalyzed removal of ubiquitin (w/o Ub), a similar immunoprecipitation and DUB protocol were used with the following modifications. The cell pellets (800 OD_600_) were divided into two equal portions. For samples processed under ‘w Ub’ condition, lysis buffer with deubiquitinase inhibitors (100 mM Tris pH 7.4, 150 mM NaCl, 5 mM EDTA, 5 mM EGTA, 10% glycerol, 0.2 % NP40, 100μM PR619, 5 mM 1,10-Phenanthroline, 50 mM N-Ethylmaleimide, phosphatase inhibitors and complete protease inhibitor tablet) was used. Immunoprecipitated samples were washed 2x. As a mock treatment for immunoprecipitated samples under ‘w Ub’ condition, the deubiquitinase buffer did not have any deubiquitinases. For the samples processed under conditions that catalyzed removal of ubiquitin (w/o Ub), the lysis buffer did not have the deubiquitinase inhibitors 100μM PR619, 5 mM 1,10- Phenanthroline, or 50 mM N-Ethylmaleimide, and furthermore the immunoprecipitated samples were processed using DUB buffer containing 1µl of each deubiquitinases Usp2, AMSH and OTULIN. Data were generated from independent experiments from three biological replicates and quantified as described later.

For systematic screening of comparative enrichment of Arf with Gos1 under various ubiquitin-preserved/ubiquitin-removed conditions in combination with phosphorylation preserved/ phosphorylation removed conditions the samples were processed as described above with the following modifications to the procedure. Cells were lysed in the presence of (1) deubiquitinase inhibitors (PR619, O-PA, NEM), (2) phosphatase inhibitors (PhosSTOP), (3) both deubiquitinase and phosphatase inhibitors or (4) no additional inhibitors other than the protease inhibitors. Cell lysis in the presence of deubiquitinase and phosphatase inhibitors is expected to preserve ubiquitin and phosphorylation-mediated complexes. Cell lysis in the presence of just deubiquitinase or phosphatase inhibitors is expected to preserve only ubiquitin or phosphorylation-mediated complexes. Cell lysis in the absence of both deubiquitinase and phosphatase inhibitors in expected to not preserve ubiquitin or phosphorylation mediated complexes. Following immunoprecipitation using anti-FLAG resin, the samples were treated with (1) deubiquitinases (USP2, AMSH and Otulin), (2) phosphatases (Lambda phosphatase), (3) both deubiquitinases and phosphatases and (4) no post-IP treatment. Post-IP deubiquitination (with USP2, AMSH and Otulin) and/or dephosphorylation (with Lambda phosphatase) is expected to strip off any preserved or remaining ubiquitination and phosphorylation, respectively, from the immunoprecipitated samples.

### Immunoblotting with ECL

Protein samples were separated by 4-20% gradient SDS-PAGE followed by immunoblotting. For anti-ubiquitin antibody the membranes were treated with glutaraldehyde solution (supplied with the antibody) as per manufacturer’s protocol and washed with PBS. The membranes were blocked in 5% non-fat milk for 1 hour, incubated with primary antibodies for 3h at room temperature, washed 5 times with TBS with 0.1% Tween, incubated with appropriate secondary antibody for 1h at room temperature, washed 5 times and imaged using manufacturer recommended chemiluminescence protocol. The membranes were imaged with AI600 Chemiluminescent Imager (GE Life Sciences). Quantitative analysis of Western Blot images was performed using ImageJ software.

### SILAC Mass spectrometry

SILAC-based mass spectrometric analysis of Gos1-FLAG with untagged control was performed using a similar protocol as described previously ^57^. Briefly, an equal amount of cells (labeled with either light or heavy Arg and Lys) expressing endogenous FLAG-tagged Gos1 or untagged cells were harvested from the mid-log phase and disrupted by bead beating using ice-cold lysis buffer (50 mM Tris-HCl, pH 7.5, 150 mM NaCl, 5 mM EDTA, 0.2% NP-40, 10 mM iodoacetamide, 1 mM 1,10-phenanthroline, 1× EDTA-free protease inhibitor cocktail [Roche], 1 mM phenylmethylsulfonyl fluoride, 20 µM MG132, 1× PhosStop [Roche], 10 mM NaF, 20 mM BGP, and 2 mM Na_3_VO_4_). Lysate was clarified by centrifugation at 21,000 × g for 10 min at 4°C and supernatant was transferred into a new tube and diluted with three-fold volume of ice-cold TBS (50 mM Tris-HCl, pH 7.5, 150 mM NaCl). Samples were incubated with 50 µL of EZview anti-FLAG M2 resin slurry (Sigma) for 2 hr at 4°C with rotation. The resin was washed three times with cold TBS and incubated with 90 µL elution buffer (100 mM Tris-HCl, pH 8.0, 1% SDS) at 98°C for 5 min. The collected eluate was reduced with 10 mM DTT, alkylated with 20 mM iodoacetamide, and precipitated with 300 µL PPT solution (50% acetone, 49.9% ethanol, and 0.1% acetic acid). Light and heavy protein pellets were dissolved with Urea-Tris solution (8 M urea, 50 mM Tris-HCl, pH 8.0). Heavy and light samples were combined, diluted four-fold with water, and digested with 1 µg MS-grade trypsin (Gold, Promega) by overnight incubation at 37°C. Phosphopeptides were enriched by immobilized metal affinity chromatography (IMAC) using Fe(III)-nitrilotriacetic acid resin as previously described (MacGurn et al., 2011) and dissolved in 0.1% trifluoroacetic acid and analyzed by LC-MS/MS using an Orbitrap XL mass spectrometer. Data collected were searched using MaxQuant (ver. 1.6.5.0) and chromatograms were visualized using Skyline (ver. 20.1.0.31, MacCoss Lab). Coimmunoprecipitation followed by SILAC-based mass spectrometric analysis of α-COP-FLAG was performed as described above.

### Statistical analysis

Statistical differences between two groups for SNARE morphology were determined using a Fisher’s exact test. For multiple group comparison, one-way ANOVA on the means using GraphPad Prism (GraphPad Software Inc.). Probability values of less than 0.05, 0.01 and 0.001 were used to show statistically significant differences and are represented with *, ** or *** respectively. To quantify Western blot data, at least three independent replicates were used, and intestines were calculated using ImageJ software and statistical analyses, as indicated, were performed using GraphPad Prism.

## Acknowledgements

We thank Dr. Scott Emr (Cornell University) and Dr. Aki Nakano (Riken Institute) for plasmids and strains. We thank Charles Barlow for the anti-COPI antibody. We thank Kristie Lindsey Rose (Vanderbilt University, Proteomics Core Laboratory) for help with processing mass spectrometry samples. These studies were supported by NIH Grants 1R01GM118452 (to TRG), 5R01GM058202 (to RCP), 1R35GM119525 (to LPJ) and 1R01GM118491 (to JAM). Lauren P Jackson is a Pew Scholar in the Biomedical Sciences, supported by the Pew Charitable Trusts.

## Author contributions

Swapneeta Date – designed and performed the majority of the experiments, analyzed results and contributed to writing - original draft, reviewing and editing. Peng Xu, Nicholas S. Diab, Jordan Best – designed and performed initial SNARE localization experiments, analyzed results, and contributed to writing - reviewing and editing. Nathaniel L. Hepowit – designed strategy and assisted with SILAC experiments and contributed to writing - reviewing and editing. Boyang Xie - designed strategy and assisted with β’COP purification and contributed to writing - reviewing and editing. Jiale Du synthesized and purified K63 ubiquitin trimers. Eric R. Strieter mentored and supervised Jiale Du and contributed to writing—reviewing and editing. Lauren P Jackson contributed to conceptualization, resources (protein purification), funding acquisition, writing—reviewing and editing, mentoring and supervision of Boyang Xie. Jason A MacGurn contributed to conceptualization, resources including DeltaVision Deconvolution Microscope, funding acquisition, writing—reviewing and editing, and mentoring and supervision of Nathaniel L. Hepowit. Todd R. Graham - vision, conceptualization, funding acquisition, methodology, project administration, writing—original draft, reviewing and editing, mentoring and supervision of Swapneeta S. Date, Peng Xu, Nicholas S. Diab and Jordan Best, and supervised the whole project.

## Competing Interests statement

All authors declare no competing financial and/or non-financial interests in relation to the work described.

## Notes

### Competing Interest Statement

The authors have declared no competing interest.

