## Supplemental Data Figures and Tables for "Ubiquitination drives COPI priming and Golgi SNARE localization"

**Extended Data for manuscript titled 'Ubiquitination regulates COPI-dependent retrieval of Golgi SNAREs'.**

#### Extended Data Figures

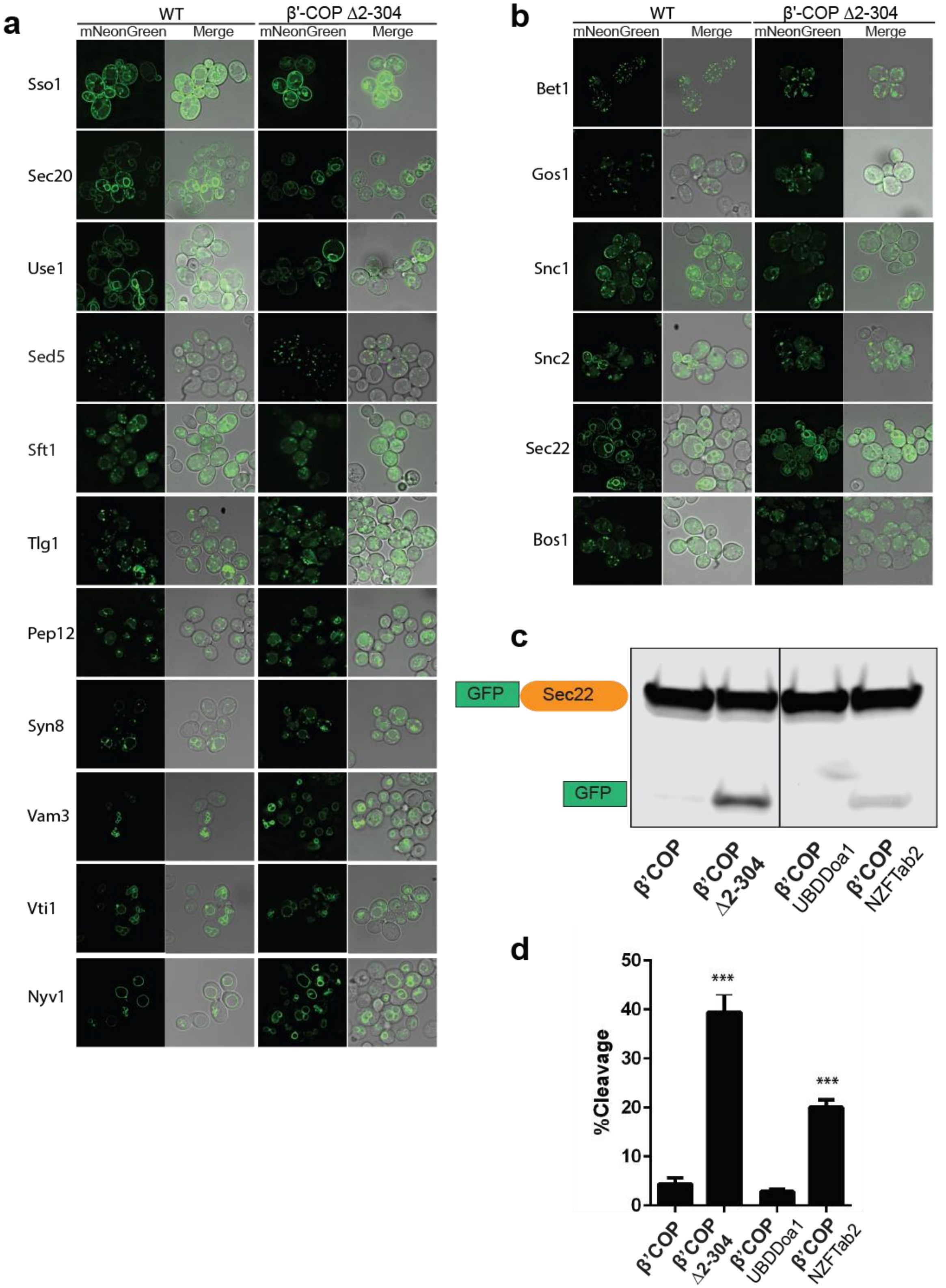

**Extended Data Figure 1. Localization of six SNAREs is perturbed in β'-COP Δ2-304 mutant:** Panels show live-cell imaging data for corresponding SNAREs tagged with mNeonGreen in *Saccharomyces cerevisiae* wild-type (WT) cells or in COPI mutant in which the N-terminal ubiquitin-binding domain of β'-COP is deleted (β'-COP Δ2-304). (**a**) No significant morphological changes are seen for eleven SNAREs in β'-COP Δ2-304 cells compared to WT cells. (**b)** Bet1 and Gos1 were observed in punctate structures in WT cells; however, in β'-COP Δ2-304 Bet1 and Gos1 were additionally seen in morphologically aberrant elongated-tube and ring-like structures. Snc1 and Snc2 are localized to plasma membrane in WT cells; however, in β'-COP Δ2-304 Snc1 and Snc2 are internalized to morphologically aberrant elongated-tube and ring-like structures. Compared to their localization in WT cells, Sec22 and Bos1 are partially mislocalized to vacuole in β'-COP Δ2-304 cells. **(c)** Western blot data using anti-GFP antibody wherein GFP-tagged Sec22 is expressed in WT cells, β'-COP Δ2-304 cells or in cells where the N-terminal WDR of β'-COP is replaced with general ubiquitin binding domain Doa1 (β'-COP UBDDoa1) or the K63-polyUb specific UBD, NZFTab2 (β'COP NZFTab2). Compared to WT cells, in β'-COP Δ2-304 GFP-Sec22 is mislocalized to vacuoles resulting in its cleavage and the release of free GFP. The mislocalization of GFP-Sec22 is cells in COPI mutant is rescued by the replacement of β'-COP-UBD with the UBD of Doa1 and partially with the UBD of Tab2. (d) Quantitation of the free GFP signal divided by the total (GFP-Sec22 + GFP) was done on three biological replicates and statistical differences were determined using a one-way ANOVA (***$p<0.001$).

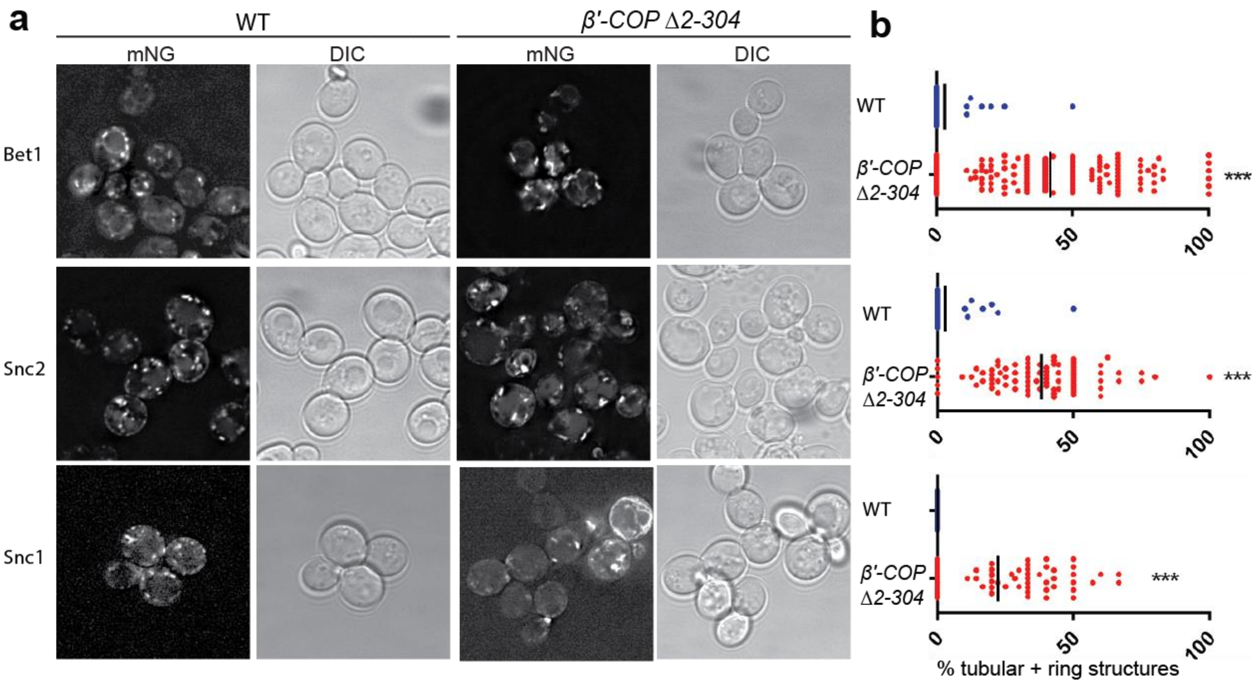

**Extended Data Figure 2. Bet1, Snc1 and Snc2 localization to aberrant membranes in β'-COP Δ2-304 mutants is independent of expression level. (a-b)** mNeonGreen (mNG) tagged SNAREs were expressed under control of the *CUP1* promoter, a weaker inducible promoter compared to constitutively expressed stronger *ADH* promoter, in WT cells or in β'-COP Δ2-304 cells. The morphological changes seen for Bet1, Snc2 and Snc1 in β'-COP Δ2-304 wherein SNAREs are mislocalized to elongated-tube and ring-like structures compared to WT cells when the SNAREs expressed under the *CUP1* promoter are similar to those observed when expressed under the *ADH* promoter. Statistical differences were determined using a one-way ANOVA on the means of the three biological replicates (***$p<0.001$).

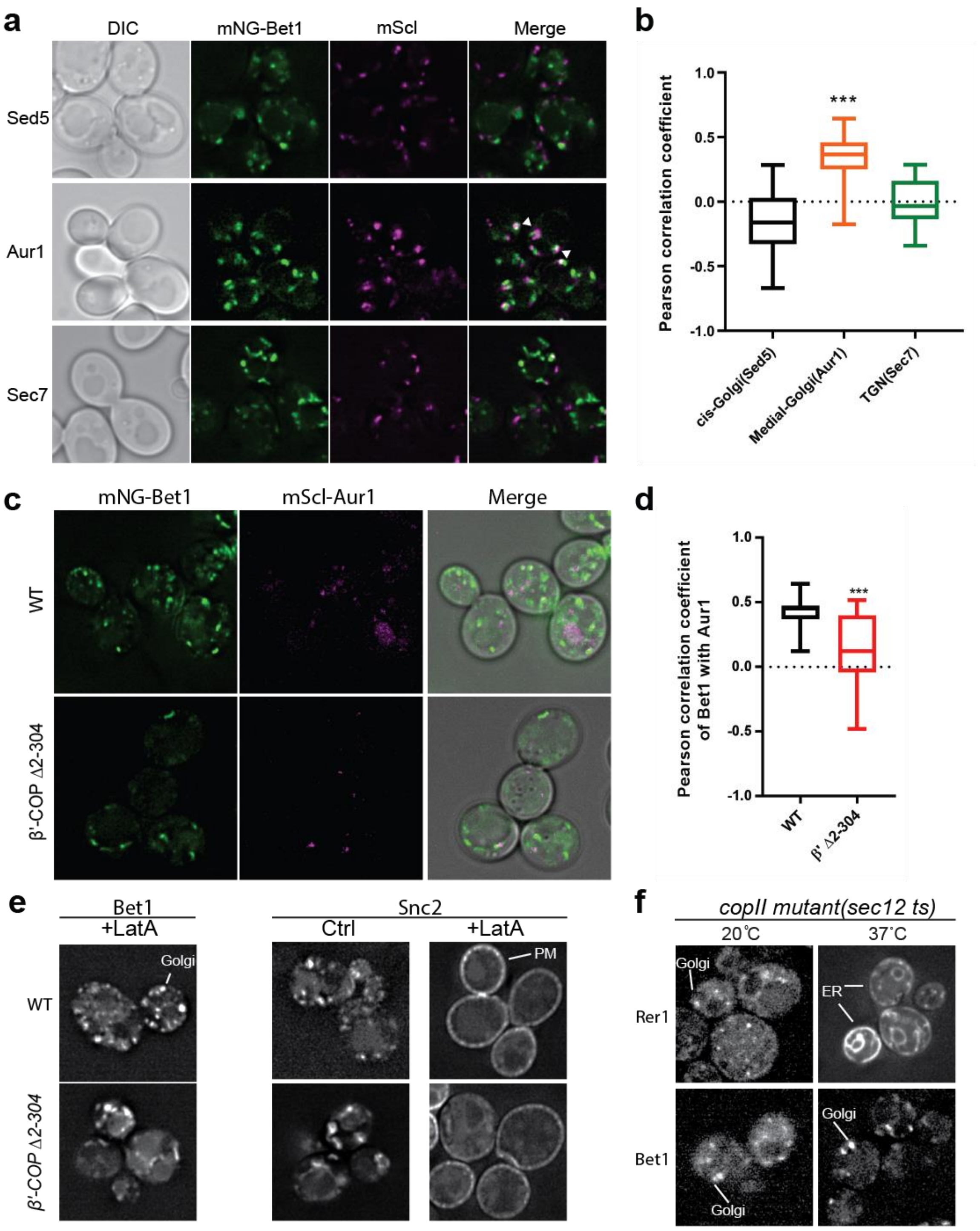

**Extended Data Figure 3. Bet1 localizes to medial Golgi and does not transit the plasma membrane or ER:** Panels a, c, e, and f show live-cell imaging data in *Saccharomyces cerevisiae*. **(a-b)** mNG-Bet1 expressed from the *CUP1* promoter colocalizes with medial Golgi marker Aur1 but not significantly with cis- and trans-Golgi network markers Sed5 and Sec7, respectively. **(c-d)** Deletion of the N-terminal WD40 propeller of β'-COP (Δ2-304) causes significantly less Bet1 colocalizing with Aur1. **(e)** Latrunculin A (LatA) treatment for 1 hour to inhibit endocytosis does not result in entrapment of Bet1 at plasma membrane (PM) in WT or β'-COP Δ2-304 cells. This is in contrast to Snc2, which is trapped at PM following LatA treatment. Thus, Snc2 cycles between the Golgi/endosomes and the plasma membrane while Bet1 follows a different trafficking itinerary. **(f)** Bet1 is localized to the Golgi in a COPII mutant (*sec12 ts*) at both permissive and non-remissive temperatures. This is in contrast to Rer1, which normally cycles between the ER and Golgi and is trapped in the ER of COPII mutants at non-permissive temperatures. These data imply that Bet1 is primarily recycled from late Golgi to early Golgi compartments by COPI.

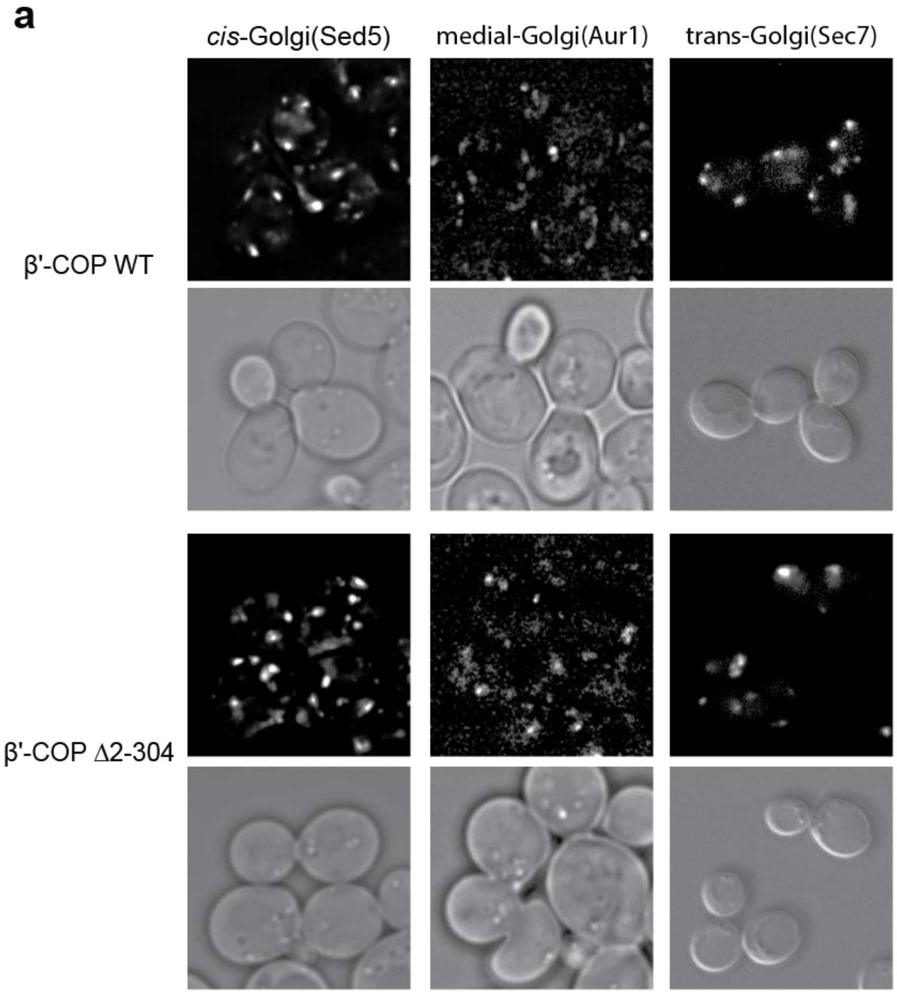

**Extended Data Figure 4. No change in *cis*, medial, and *trans*-Golgi network morphology in β'-COP Δ2-304 cells.** No apparent changes are observed in the morphology of Golgi membranes containing *cis*-, medial- and *trans*-Golgi network markers, Sed5, Aur1 and Sec7, respectively, in β'-COP Δ2-304 cells compared to WT cells.

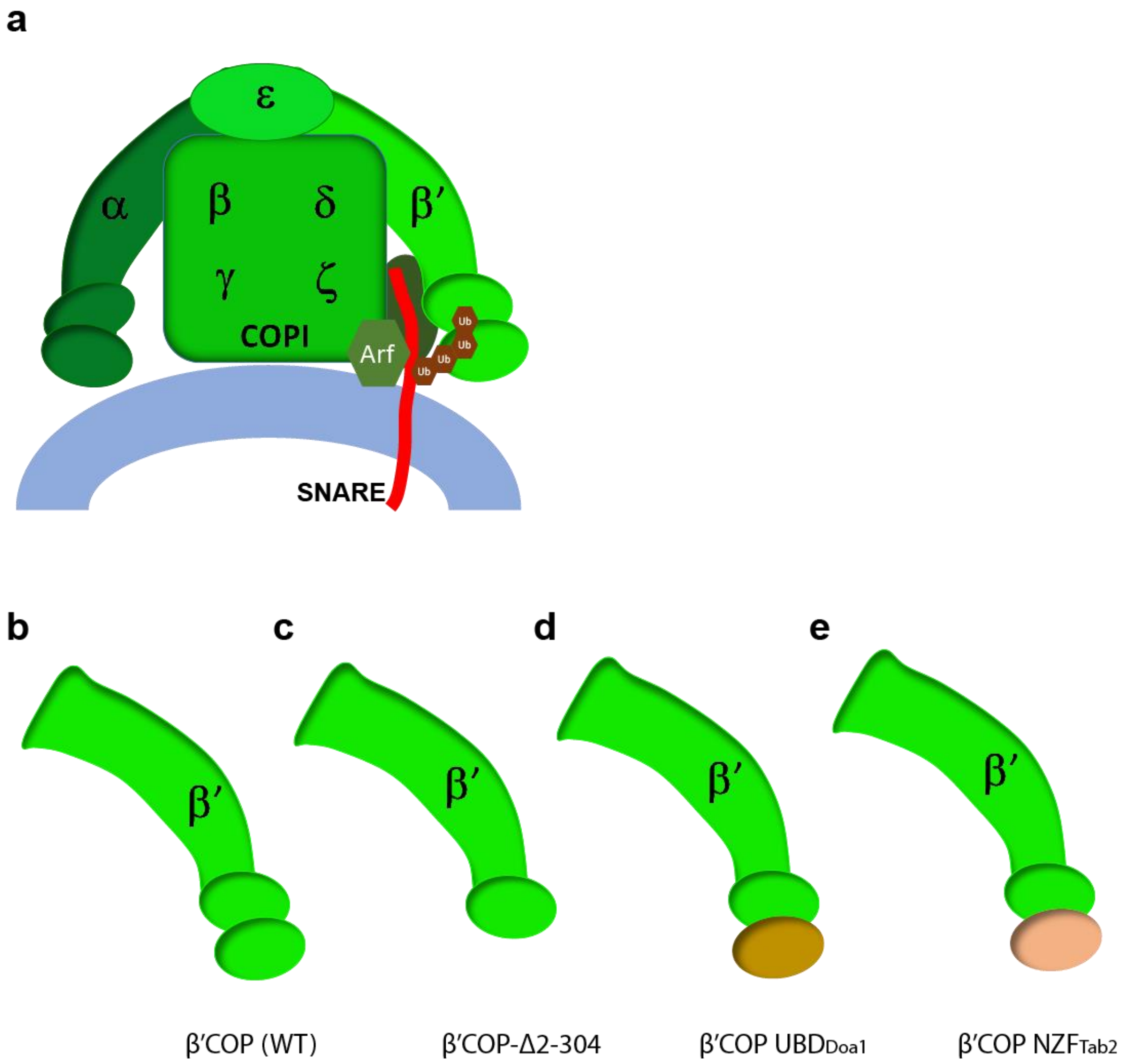

**Extended Data Figure 5. COPI Model figure with β'-COP constructs that differ in their ability to recognize and bind ubiquitin linkages. (a)** COPI model figure depicting all COPI subunits, the N-terminal WDR of β'-COP that binds ubiquitin, key components of COPI coat complex including Arf, ArfGAP (dark oval behind the SNARE) and a SNARE (red). **(b-e)** β'-COP constructs that differ in their ability to recognize and bind polyubiquitin linkages. The N-terminal WDR of β'-COP **(b)** is deleted to generate β'-COP Δ2-304 **(c)** or replaced with a general ubiquitin-binding domain from Doa1 that can bind any polyubiquitin linkage (β'-COP UBDDoa1) **(d)** or the K63-polyUb specific UBD, NZFTab2 (β'COP NZFTab2) **(e)**.

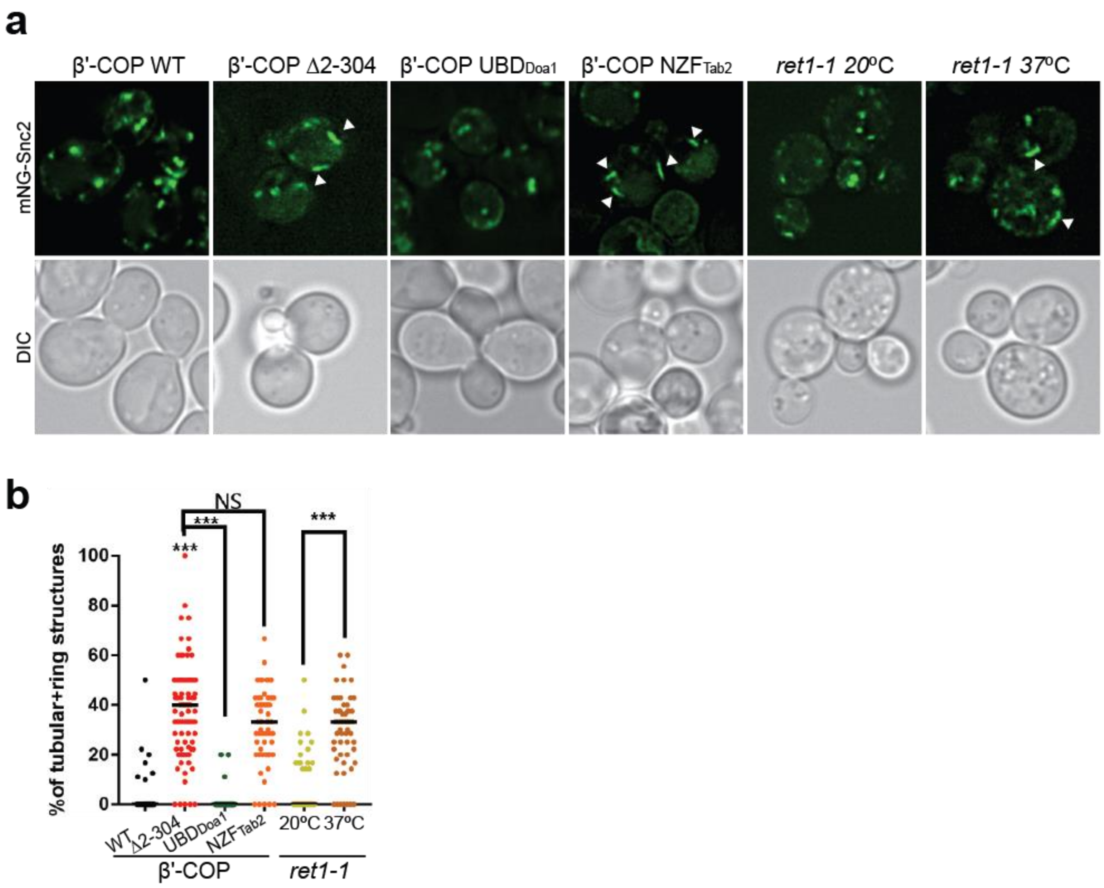

**Extended Data Figure 6. The ability of β'COP to bind ubiquitin is critical for the proper localization of Snc2. (a-b)** Deletion of the N-terminal ubiquitin-binding WDR of β'COP leads to accumulation of mNG-Snc2 into elongated tubular and ring-like structures (white arrowheads). This phenotype is rescued by the replacement of the N-terminal ubiquitin-binding WD4R of β'COP by the general ubiquitin bunding domain Doa1 (β'-COP UBD$_{Doa1}$) but not when replaced with the K63-specific UBD, NZF$_{Tab2}$ (β'COP NZF$_{Tab2}$). The mislocalization of Snc2 observed in Δ2-304 cells is comparable to COPI inactivation phenotype observed for *ret1-1* at nonpermissive temperatures. Statistical differences were determined using a one-way ANOVA on the means of the three biological replicates (***p<0.001).

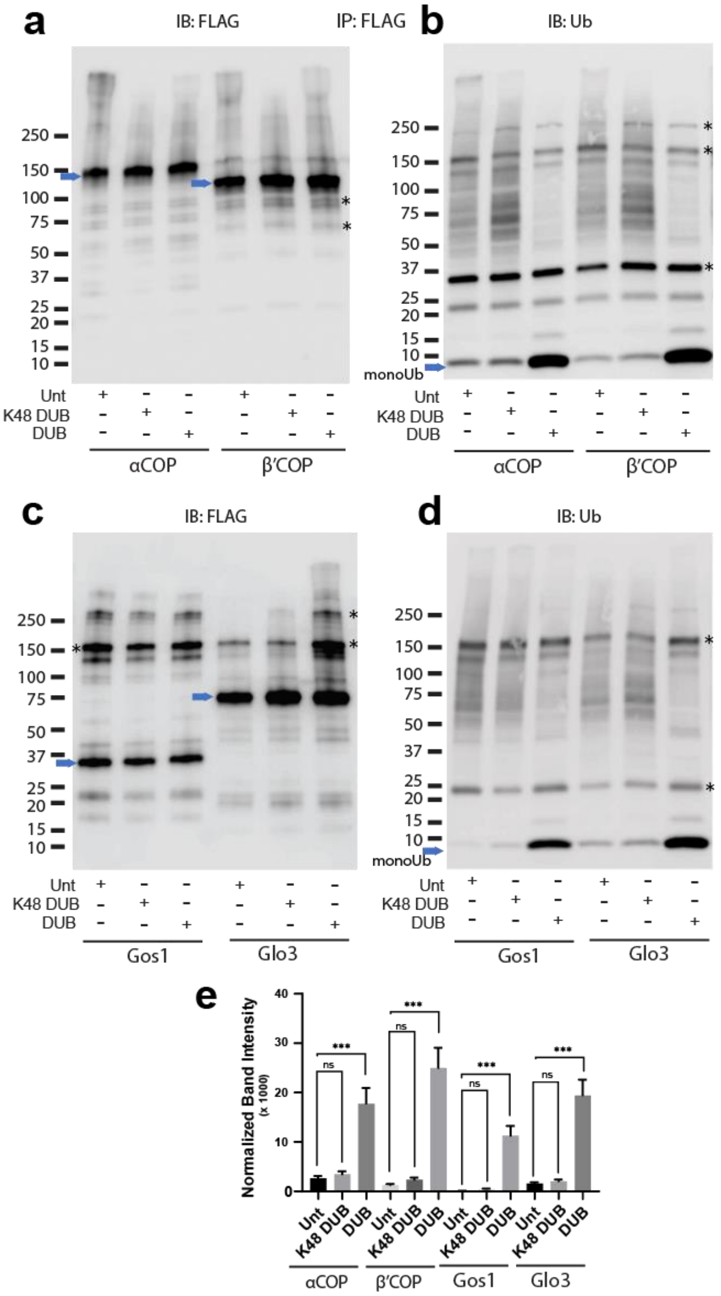

**Extended Data Figure 7 Ubiquitination associated with COPI, Gos1, and Glo3 is non-K48 linked.** (a-d) Western Blot data wherein - FLAG-tagged α-COP or β'COP, Gos1, and Glo3 were immunoprecipitated and treated with a general deubiquitinase (DUB), a K48-specific deubiquitinases (K48-DUB) or mock-treated (Unt) and probed with (a and c) FLAG or (b and d) ubiquitin antibody. (b and d) No significant release of ubiquitin is observed following the K48-DUB treatment. Blue arrows indicate the position of the FLAG-tagged protein and the asterisks indicate the position of prominent background bands. (e) Statistical differences were determined using a one-way ANOVA with multiple comparison test on the means of the three biological replicates (***p<0.001, NS p > 0.05).

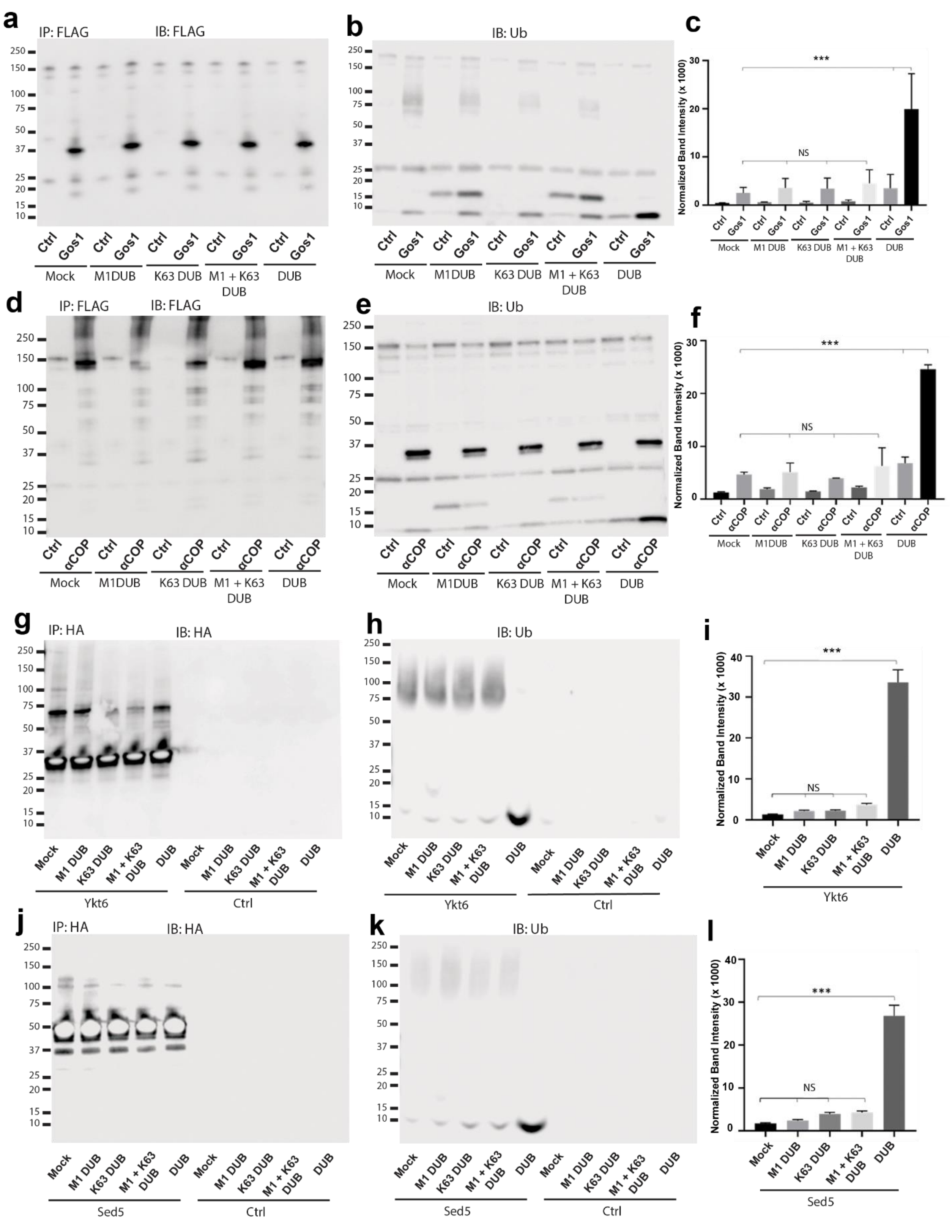

**Extended Data Figure 8. Ubiquitin associated with Gos1, α-COP, Ykt6 and Sed5 is not K63- or M1-linked polyubiquitin.** Western blot analysis for FLAG-tagged Gos1 and α-COP, and HA-tagged Ykt6 and Sed5 or untagged cells (Ctrl) following immunoprecipitation, treatment with mock buffer (no DUB), linkage specific deubiquitinases (K63 DUB, M1 DUB or M1 + K63

DUB) or with general deubiquitinase (DUB). Western blots were probed with with (**a,d**) FLAG or (**g, j**) HA antibody and (**b, e, h, k**) ubiquitin antibodies. Blots probed with FLAG or HA antibody show equal amount of corresponding bait recovered under different conditions. The treatment of immunoprecipitated samples with general deubiquitinases leads to significant release of monoubiquitin from Gos1, α-COP, Ykt6 and Sed5 but no significant release of monoubiquitin is observed when treated with linkage-specific K63-DUB, M1-DUB or the K63+M1-DUB combination. Statistical differences were determined using a one-way ANOVA with multiple comparison test on three biological replicates (****$p \leq 0.0001$, ***$p<0.001$, *$p<0.05$, Ns $p > 0.05$).

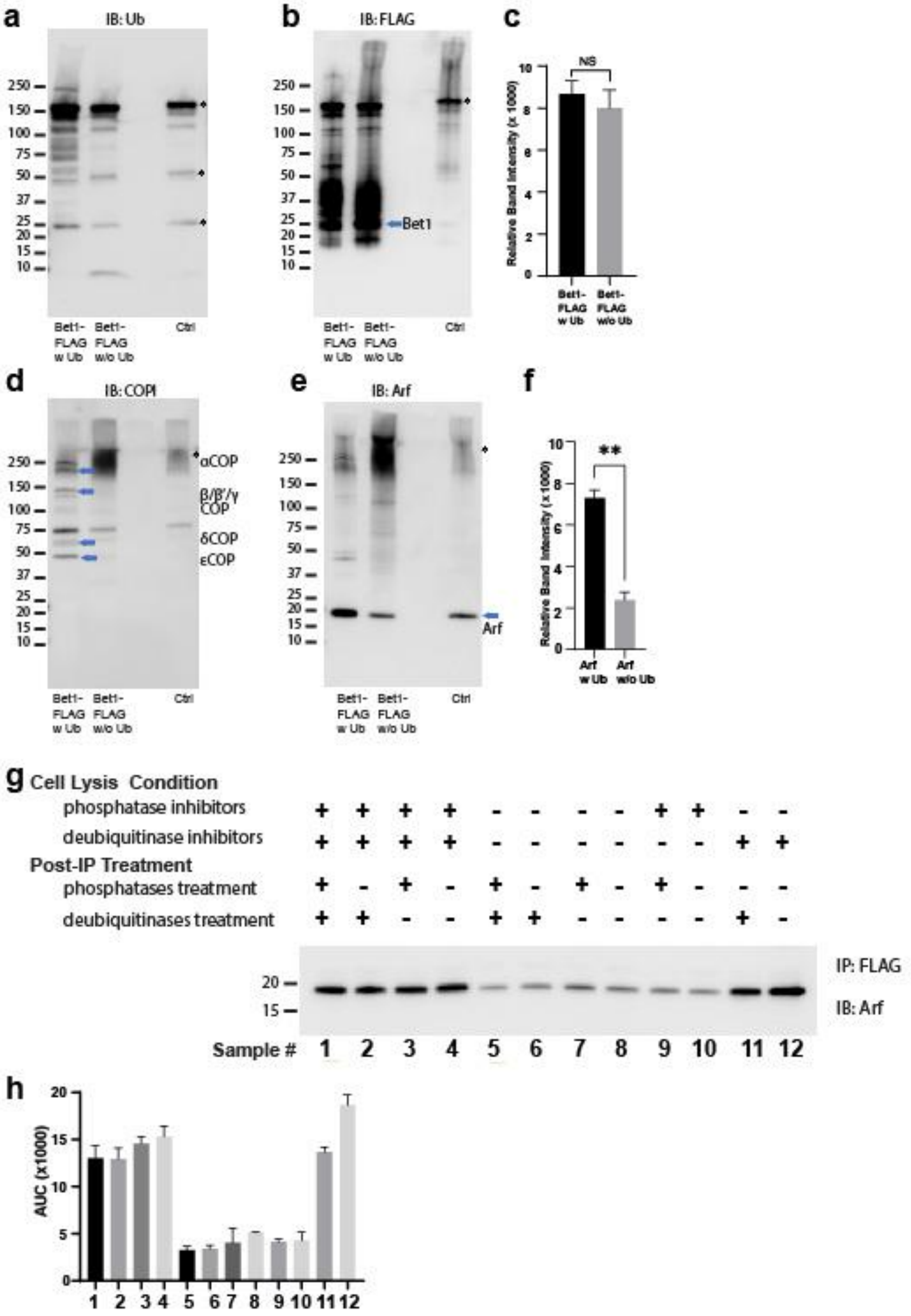

**Extended Data Figure 9. Ubiquitin-mediated enhancement of COPI and Arf Bet1:** Western Blot data showing comparative pulldowns of Bet1-FLAG **(a-f)** processed under 'ubiquitin-preserved' (w Ub) and 'no-ubiquitin' (w/o Ub) condition, and probed for Ub **(a)**, FLAG **(b)**, COPI **(d)** and Arf **(e)**. Untagged cells processed under 'w UB' condition to determine background binding were used as a control (Cntr) and abundant background bands are marked with an asterisk. Quantitation of Bet1-FLAG **(c)** and Arf **(f)** in the pulldown samples. Band intensities are measured using ImageJ. Quantitation was done on three biological replicates using a t-test ***p<0.001, Ns p > 0.05). **(g)** Western Blot data for Arf enriched with Gos1-FLAG under various combinations of cell lysis conditions to either preserve or not preserve ubiquitination and/or phosphorylation and to strip off ubiquitin and/or phosphorylation by post-immunoprecipitation (post-IP) treatments as indicated. Cell lysis in the presence of deubiquitinase and phosphatase inhibitors is expected to preserve ubiquitin and phosphorylation-mediated complexes. Cell lysis in the presence of just deubiquitinase or phosphatase inhibitors is expected to preserve only ubiquitin or phosphorylation-mediated complexes. Cell lysis in the absence of both deubiquitinase and phosphatase inhibitors in expected to not preserve ubiquitin or phosphorylation mediated complexes. Post-IP deubiquitination (with USP2, AMSH and Otulin) and/or dephosphorylation (with Lambda phosphatase) is expected to strip off any preserved or remaining ubiquitination and phosphorylation, respectively, from the immunoprecipitated samples. Band intensities were measured using ImageJ and quantitation was done on three biological replicates.

### Extended Data Tables

| SNARE Name | Classification | Localization |
|---|---|---|
| Bet1 | Qc | Golgi |
| Gos1 | Qb | ER-Golgi |
| Snc1 | R | PM-vesicles |
| Snc2 | R | PM-vesicles |
| Sec22 | R | ER-Golgi |
| Bos1 | Qb | ER-Golgi |
| SSo1 | Qa | PM |
| Sec20 | Qb | ER |
| Use1 | Qc | ER |
| Sed5 | Qa | Golgi |
| Sft1 | Qc | Golgi |
| Tlg1 | Qc | Late Golgi |
| Pep12 | Qa | Golgi-Vacuole |
| Syn8 | Qc | Golgi-Vacuole |
| Vam3 | Qa | Vacuole |
| Vti1 | Qb | Golgi-Vacuole |
| Nyv1 | R | Golgi-Vacuole |

**Table 1: *Saccharomyces cerevisiae* SNAREs used in this study.**

| Name | Genotype | Sources | Additional strains |
|---|---|---|---|
| BY4742 | *MATα his3 leu2 ura3 lys2* | Invitrogen | |
| β'COP WT | *MATα his3 leu2 ura3 lys2 sec27Δ::Hygro p315-SEC27* | Xu et al., 2017 | Additional strains were generated by transformaring this strain with different plasmids encoding mNG-tagged SNAREs. Plasmids are listed in the 'Plasmids used in this study' table. Some strains were generated by transforming two plasmids with different selection markers. |
| β'COP-Δ2-304 | *MATα his3 leu2 ura3 lys2 sec27Δ::Hygro p315-sec27Δ2-304* | Xu et al., 2018 | Additional strains were generated by transformaring this strain with different plasmids encoding mNG-tagged SNAREs. Plasmids are listed in the 'Plasmids used in this study' table. |
| β'COP-UBDDoa1 | *MATα his3 leu2 ura3 lys2 sec27Δ::Hygro p315-DOA1(1-450)-sec27(305-889)* | Xu et al., 2019 | Additional strains were generated by transformaring this strain with different plasmids encoding mNG-tagged SNAREs. Plasmids are listed in the 'Plasmids used in this study' table. Some strains were generated by transforming two plasmids with different selection markers. |
| β'COP-NZFTab2 | *MATα his3 leu2 ura3 lys2 sec27Δ::Hygro p315-NZF-sec27(305-899)* | Xu et al., 2020 | Additional strains were generated by transformaring this strain with different plasmids encoding mNG-tagged SNAREs. Plasmids are listed in the 'Plasmids used in this study' table. |
| Bet1-HTF | *MATα his3 leu2 ura3 lys2 BET1::6xHIS-TEV-3xFLAG::ClonNAT* | This study | |
| Gos1-HTF | *MATα his3 leu2 ura3 lys2 GOS1::UL36*-3xHA::ClonNAT* | This study | |
| Snc1-HTF | *MATα his3 leu2 ura3 lys2 SNC1::UL36-3xHA::ClonNAT* | This study | |
| Art1-HTF | *MATα his3 leu2 ura3 lys2 pRS416-pART1-ART1-3xFLAG* | This study | |
| SILACα | *MATα Δarg4::kanMX6 leu2-3,112 ura3-52 his3-Δ200 trp1-Δ901 suc2-Δ9 lys2-801* | Lee et al., 2017 | |
| SILAC-Gos1-HTF | *MATa Δarg4::kanMX6 leu2-3,112 ura3-52 his3-Δ200 trp1-Δ901 suc2-Δ9 lys2-801 GOS1::6xHIS-TEV-3xFLAG::ClonNAT* | This study | |
| α-COP-DUB | *MATα his3 leu2 ura3 lys2 SEC27::UL36-3xHA::ClonNAT* | This study | |
| α-COP-DUB* | *MATα his3 leu2 ura3 lys2 SEC27::UL36*-3xHA::ClonNAT* | This study | |
| β'-COP-DUB | *MATα his3 leu2 ura3 lys2 COP1::UL36-3xHA::ClonNAT* | This study | |
| β'-COP-DUB* | *MATα his3 leu2 ura3 lys2 COP1::UL36*-3xHA::ClonNAT* | This study | |
| 3xHA Ykt6 | *MATα his3 leu2 ura3 lys2 ::ClonNAT-TEF-3xHA::YKT6* | This study | |
| 3xHA Sed5 | *MATα his3 leu2 ura3 lys2 ::ClonNAT-TEF-3xHA::SED5* | This study | |
| α-COP-HTF | *MATα his3 leu2 ura3 lys2 SEC27::6xHIS-TEV-3xFLAG::ClonNAT* | This study | |
| β'-COP-HTF | *MATα his3 leu2 ura3 lys2 COP1::6xHIS-TEV-3xFLAG::ClonNAT* | This study | |
| Glo3-HTF | *MATα his3 leu2 ura3 lys2 GLO3::6xHIS-TEV-3xFLAG::ClonNAT* | This study | |
| EGY101-16d | *MATa ret1-1 leu2-3,112 ura3-52 his3-Δ200 trp1-Δ901 suc2-Δ9* | (Gaynor and Emr, 1997) | |

| | | |
|---|---|---|
| PXY2103A | *BY4742 p416-GFP-Rer1 p313-mCherry-Tlg1* | This study |

**Table 2: List of strains used in this study.**

| **Name** | **Plasmids** | **Description** | **Sources** |
|---|---|---|---|
| Vector | pRS416 | Yeast shuttle vector | ATCC |
| β'-COP WT | pRS315-SEC27 | *SEC27* (b'-COP) whole coding cassette including its promoter and terminator | Xu et al., 2017 |
| β'-COP Δ2-304 | pRS315-sec27Δ2-304 | *SEC27* with deletion the N-terminal b-propeller | Xu et al., 2017 |
| ADH-mNG-Bet1 | pRS416-ADH-mNG-Bet1 | *mNG tagged Bet1 under ADH promoter* | This study |
| mNG-Bet1 | pRS416-Cup1-mNG-Bet1 | *mNG tagged SNARE under CUP1 promoter* | This study |
| mNG-Gos1 | pRS416-Cup1-mNG-Gos1 | *mNG tagged SNARE under CUP1 promoter* | This study |
| mNG-Snc2 | pRS416-Cup1-mNG-Snc2 | *mNG tagged SNARE under CUP1 promoter* | This study |
| mNG-Snc1 | pRS416-Cup1-mNG-Snc1 | *mNG tagged SNARE under CUP1 promoter* | This study |
| mNG-Sec22 | pRS416-Cup1-mNG-Sec22 | *mNG tagged SNARE under CUP1 promoter* | This study |
| mNG-Bos1 | pRS416-Cup1-mNG-Bos1 | *mNG tagged SNARE under CUP1 promoter* | This study |
| mNG-SSo1 | pRS416-Cup1-mNG-SSo1 | *mNG tagged SNARE under CUP1 promoter* | This study |
| mNG-Sec20 | pRS416-Cup1-mNG-Sec20 | *mNG tagged SNARE under CUP1 promoter* | This study |
| mNG-Use1 | pRS416-Cup1-mNG-Use1 | *mNG tagged SNARE under CUP1 promoter* | This study |
| mNG-Sed5 | pRS416-Cup1-mNG-Sed5 | *mNG tagged SNARE under CUP1 promoter* | This study |
| mNG-Sft1 | pRS416-Cup1-mNG-Sft1 | *mNG tagged SNARE under CUP1 promoter* | This study |
| mNG-Tlg1 | pRS416-Cup1-mNG-Tlg1 | *mNG tagged SNARE under CUP1 promoter* | This study |
| mNG-Pep12 | pRS416-Cup1-mNG-Pep12 | *mNG tagged SNARE under CUP1 promoter* | This study |
| mNG-Syn8 | pRS416-Cup1-mNG-Syn8 | *mNG tagged SNARE under CUP1 promoter* | This study |
| mNG-Vam3 | pRS416-Cup1-mNG-Vam3 | *mNG tagged SNARE under CUP1 promoter* | This study |
| mNG-Vti1 | pRS416-Cup1-mNG-Vti1 | *mNG tagged SNARE under CUP1 promoter* | This study |
| mNG-Nyv1 | pRS416-Cup1-mNG-Nyv1 | *mNG tagged SNARE under CUP1 promoter* | This study |
| mScl-Sed5 | pRS315-ADH-mScl-Sed5 | *mNG tagged SNARE under CUP1 promoter* | This study |
| mScl-Aur1 | pRS315-ADH-mScl-Aur1 | *mNG tagged SNARE under CUP1 promoter* | This study |
| mScl-Sec7 | pRS315-ADH-mScl-Sec7 | *mNG tagged SNARE under CUP1 promoter* | This study |
| β'-COP UBD$_{Doa1}$ | pRS315-Doa1(1-450)-Sec27(305-889) | *SEC27* first b-propeller replaced with DOA1(1-450) | Xu et al., 2017 |
| β'-COP NZF$_{Tab2}$ | pRS315-NZF-Sec27(305-899) | *SEC27* first b-propeller replaced with NZF (TAB2 aa 665-693) | Xu et al., 2017 |
| human beta-Propeller-GST | pCOPB1(1-604)-GST | human b'-COP N-terminal propeller tagged with GST | Xu et al., 2017 |

**Table 3: List of plasmids used in this study.**

| Protein IDs | Log2 of Normalized H/L Values |
|---|---|
| sp\|P38736\|GOSR1_YEAST | -4.9059964018546 |
| sp\|P43682\|SFT1_YEAST | -3.97068423418304 |
| sp\|P40509\|COPE_YEAST | -3.49365398192928 |
| sp\|P32602\|SEC17_YEAST | -3.33741894711022 |
| sp\|Q01590\|SED5_YEAST | -2.82500562887929 |
| sp\|P22213\|SLY1_YEAST | -2.71151149773846 |
| sp\|Q03532\|HAS1_YEAST | -2.62941162387656 |
| sp\|P00359\|G3P3_YEAST | -1.97573706508829 |
| sp\|P00358\|G3P2_YEAST | -1.81975027517981 |
| sp\|P07253\|CBP6_YEAST | -1.72690191082151 |
| sp\|P05318\|RLA1_YEAST | -1.69282797672698 |
| sp\|P32582\|CBS_YEAST | -1.65140496929358 |
| sp\|P53622\|COPA_YEAST | -1.64968380836001 |
| sp\|P40008\|FMP52_YEAST | -1.593920536838 |
| sp\|P22354\|RM20_YEAST | -1.57841312683729 |
| sp\|P22353\|RM08_YEAST | -1.55152187858852 |
| sp\|P41810\|COPB_YEAST | -1.45707833932508 |
| sp\|P41811\|COPB2_YEAST | -1.45225406131286 |
| sp\|P38701\|RS20_YEAST | -1.39175848673728 |
| sp\|P0CX54\|RL12B_YEAST | -1.35228946572746 |
| sp\|P41940\|MPG1_YEAST | -1.32225273779526 |
| sp\|P28241\|IDH2_YEAST | -1.23912333530525 |
| sp\|P36015\|YKT6_YEAST | -1.1169110067272 |
| sp\|P39990\|SNU13_YEAST | -1.0144121362815 |
| sp\|P00950\|PMG1_YEAST | -0.999798036831368 |
| sp\|P32316\|ACH1_YEAST | -0.967447723886688 |
| sp\|P12695\|ODP2_YEAST | -0.906899612480702 |
| sp\|Q3E7X9\|RS28A_YEAST | -0.892044369912607 |
| sp\|P54115\|ALDH6_YEAST | -0.841740539167545 |
| sp\|P43588\|RPN11_YEAST | -0.828357611061281 |
| sp\|Q3E7Y3\|RS22B_YEAST | -0.818757481214114 |
| sp\|P00330\|ADH1_YEAST | -0.810864937404904 |
| sp\|P00549\|KPYK1_YEAST | -0.807098866412113 |
| sp\|P15180\|SYKC_YEAST | -0.805509453253494 |
| sp\|P19882\|HSP60_YEAST | -0.792132130307272 |
| sp\|P12686\|RT13_YEAST | -0.791582627341841 |
| sp\|P0CS90\|HSP77_YEAST | -0.771987905289947 |
| sp\|P14540\|ALF_YEAST | -0.754114965508701 |
| sp\|P0CX42\|RL23B_YEAST | -0.710354366162678 |
| sp\|P07991\|OAT_YEAST | -0.708702929351299 |
| sp\|P0CX52\|RS16B_YEAST | -0.701060210424078 |

| | |
|---|---|
| sp\|P25694\|CDC48_YEAST | -0.686080854141113 |
| sp\|P33299\|PRS7_YEAST | -0.682788563506934 |
| sp\|P39954\|SAHH_YEAST | -0.681353437479513 |
| sp\|P05317\|RLA0_YEAST | -0.675143323222093 |
| sp\|P26783\|RS5_YEAST | -0.674982078057249 |
| sp\|Q03940\|RUVB1_YEAST | -0.672450589355627 |
| sp\|P07703\|RPAC1_YEAST | -0.666599166377839 |
| sp\|P25605\|ILV6_YEAST | -0.639062692603436 |
| sp\|P00925\|ENO2_YEAST | -0.634083546141939 |
| sp\|P15108\|HSC82_YEAST | -0.627471303297207 |
| sp\|P14120\|RL30_YEAST | -0.619536510579312 |
| sp\|P53032\|SUT1_YEAST | -0.599330922282834 |
| sp\|P02294\|H2B2_YEAST | -0.587966431578442 |
| sp\|P05750\|RS3_YEAST | -0.563134578598506 |
| sp\|P32481\|IF2G_YEAST | -0.554103827010342 |
| sp\|P38708\|YHI0_YEAST | -0.553637888121405 |
| sp\|P05738\|RL9A_YEAST | -0.548290378898981 |
| sp\|P33298\|PRS6B_YEAST | -0.542836506093522 |
| sp\|P38011\|GBLP_YEAST | -0.538932506073829 |
| sp\|Q12118\|SGT2_YEAST | -0.538408579367623 |
| sp\|P53090\|ARO8_YEAST | -0.51814310380771 |
| sp\|P39567\|IMDH1_YEAST | -0.517234306489234 |
| sp\|P47169\|MET5_YEAST | -0.513090020563888 |
| sp\|P06169\|PDC1_YEAST | -0.51179351192034 |
| sp\|O13516\|RS9A_YEAST | -0.510210470896386 |
| sp\|Q12154\|GET3_YEAST | -0.502382525244903 |
| sp\|P47079\|TCPQ_YEAST | -0.479733658969423 |
| sp\|P39076\|TCPB_YEAST | -0.478768309216214 |
| sp\|Q12251\|YP011_YEAST | -0.477562529674889 |
| sp\|P43616\|DUG1_YEAST | -0.470668778534137 |
| sp\|P05755\|RS9B_YEAST | -0.462753639020927 |
| sp\|P0CX36\|RS4B_YEAST | -0.461720104381538 |
| sp\|P32835\|GSP1_YEAST | -0.460508631722068 |
| sp\|P07244\|PUR2_YEAST | -0.454901462675821 |
| sp\|P35194\|UTP20_YEAST | -0.449276605701977 |
| sp\|P39078\|TCPD_YEAST | -0.448488904154508 |
| sp\|Q02948\|BECN1_YEAST | -0.44701312195081 |
| sp\|Q01939\|PRS8_YEAST | -0.439206799780796 |
| sp\|P02994\|EF1A_YEAST | -0.435358438634197 |
| sp\|Q03558\|OYE2_YEAST | -0.426916368814823 |
| sp\|P25367\|RNQ1_YEAST | -0.404973815934146 |
| sp\|P00958\|SYMC_YEAST | -0.403522777299489 |
| sp\|P05756\|RS13_YEAST | -0.399520899603532 |

| | |
|---|---|
| sp\|P53549\|PRS10_YEAST | -0.398493637076836 |
| sp\|P40319\|ELO3_YEAST | -0.38243820702281 |
| sp\|P18759\|SEC18_YEAST | -0.377613104102919 |
| sp\|P39676\|FHP_YEAST | -0.374916582077966 |
| sp\|P06787\|CALM_YEAST | -0.371926345803864 |
| sp\|P38711\|RS27B_YEAST | -0.365462488961771 |
| sp\|Q00055\|GPD1_YEAST | -0.363196757319526 |
| sp\|P33297\|PRS6A_YEAST | -0.358934949426979 |
| sp\|P20606\|SAR1_YEAST | -0.357843729014171 |
| sp\|P40413\|TCPE_YEAST | -0.356310018267753 |
| sp\|P33892\|GCN1_YEAST | -0.354095483897873 |
| sp\|Q04062\|RPN9_YEAST | -0.349327335213578 |
| sp\|P32939\|YPT7_YEAST | -0.345747786563309 |
| sp\|P17255\|VATA_YEAST | -0.34391555995066 |
| sp\|P25443\|RS2_YEAST | -0.343677541313105 |
| sp\|P53128\|MTHR2_YEAST | -0.340513795287394 |
| sp\|P05744\|RL33A_YEAST | -0.32908716870168 |
| sp\|P39077\|TCPG_YEAST | -0.327656130166519 |
| sp\|P40150\|HSP76_YEAST | -0.322703751948355 |
| sp\|P36150\|SUMT_YEAST | -0.311452578746133 |
| sp\|P40341\|YTA12_YEAST | -0.308644508068939 |
| sp\|P41056\|RL33B_YEAST | -0.30721574336599 |
| sp\|P36105\|RL14A_YEAST | -0.306626789898058 |
| sp\|P38230\|QOR_YEAST | -0.297785748989559 |
| sp\|P11746\|MCM1_YEAST | -0.295676895103526 |
| sp\|Q06668\|OMS1_YEAST | -0.295198844399873 |
| sp\|P33201\|MRT4_YEAST | -0.289333504176652 |
| sp\|P07342\|ILVB_YEAST | -0.285002827960929 |
| sp\|P60010\|ACT_YEAST | -0.284897363942395 |
| sp\|Q06697\|CDC73_YEAST | -0.276240501811943 |
| sp\|P42943\|TCPH_YEAST | -0.275803779390026 |
| sp\|P10659\|METK1_YEAST | -0.274948186742953 |
| sp\|P16140\|VATB_YEAST | -0.274162885112468 |
| sp\|P02557\|TBB_YEAST | -0.272977017520218 |
| sp\|P36008\|EF1G2_YEAST | -0.267583173344987 |
| sp\|P04840\|VDAC1_YEAST | -0.266958096351829 |
| sp\|P14020\|DPM1_YEAST | -0.26321331111122 |
| sp\|P09064\|IF2B_YEAST | -0.26231323859014 |
| sp\|P38077\|ATPG_YEAST | -0.261327265333245 |
| sp\|P32905\|RSSA1_YEAST | -0.26079131702105 |
| sp\|P16521\|EF3A_YEAST | -0.260549341094133 |
| sp\|P19358\|METK2_YEAST | -0.259599114098972 |
| sp\|P53252\|PIL1_YEAST | -0.242345179134673 |

| | |
|---|---|
| sp\|P32476\|ERG1_YEAST | -0.240537333120423 |
| sp\|Q04373\|PUF6_YEAST | -0.240111284295665 |
| sp\|P53981\|YNB0_YEAST | -0.239719430468653 |
| sp\|P02309\|H4_YEAST | -0.237982925406972 |
| sp\|P16387\|ODPA_YEAST | -0.236078579238591 |
| sp\|Q3E754\|RS21B_YEAST | -0.23212489511273 |
| sp\|P38431\|IF5_YEAST | -0.230397501766746 |
| sp\|P06634\|DED1_YEAST | -0.229111767692722 |
| sp\|P39079\|TCPZ_YEAST | -0.227252863891037 |
| sp\|P07251\|ATPA_YEAST | -0.222111123229149 |
| sp\|P26785\|RL16B_YEAST | -0.220984076809529 |
| sp\|Q03529\|SCS7_YEAST | -0.220009135176291 |
| sp\|P12612\|TCPA_YEAST | -0.208044270431714 |
| sp\|P00830\|ATPB_YEAST | -0.207028070930172 |
| sp\|P05030\|PMA1_YEAST | -0.205363717084051 |
| sp\|P11076\|ARF1_YEAST | -0.205330449584 |
| sp\|P07283\|PMM_YEAST | -0.195467274839513 |
| sp\|P26786\|RS7A_YEAST | -0.191623187454742 |
| sp\|P08456\|PSS_YEAST | -0.190470309408274 |
| sp\|P14742\|GFA1_YEAST | -0.190470309408274 |
| sp\|O14455\|RL36B_YEAST | -0.186869386492871 |
| sp\|Q12213\|RL7B_YEAST | -0.186688755060131 |

**Table 4: SILAC top 150 hits based on normalized H/L ratio**
